# The skin environment controls local dendritic cell differentiation and function through innate IL-13

**DOI:** 10.1101/2021.01.05.425466

**Authors:** Johannes U Mayer, Olivier Lamiable, Kerry L Hilligan, Jodie S Chandler, Samuel I Old, David A Eccles, Jianping Yang, Greta R Webb, Rita G Domingues, Luis Munoz-Erazo, Kirsty A Wakelin, Evelyn J Hyde, Shiau-Choot Tang, Sally C Chappell, Charles R Mackay, Frank Brombacher, Alan Sher, Roxane Tussiwand, Lisa M Connor, Dragana Jankovic, Matthew R Hepworth, Graham Le Gros, Franca Ronchese

## Abstract

The signals driving the adaptation of type-2 dendritic cells (DC2s) to diverse peripheral environments are not well understood. We show that the development of CD11b^low^ migratory DC2s, a DC2 population unique to the dermis, requires STAT6- and KLF4-dependent IL-13 signaling, whereas DC2s in lung and small intestine are STAT6-independent. Dermal IL-13 is mostly derived from innate lymphoid cells expressing a resting ICOS+ KLRG1-ST2-phenotype. Analysis of public datasets indicates that human skin DC2s also express an IL-4/IL-13 gene signature compared to blood or spleen, suggesting a similar developmental pathway in mice and humans. In the absence of IL-13 signaling, dermal DC2s are stable in number but remain CD11b^hi^ and show defective activation in response to allergen with diminished ability to support IL-4+ GATA3+ Th development, whereas anti-fungal IL-17+ ROR*γ*t+ responses are increased. Thus, steady-state IL-13 fosters a non-inflammatory and pro-allergic environment in healthy skin via conditioning of local DC2s.

## INTRODUCTION

DCs play a crucial role in informing the adaptive immune system about infectious, innocuous or self antigens and directing T cell responses towards effector activation, ignorance or tolerance (Eisenbarth, 2019). Two developmentally distinct lineages of DCs termed ‘type 1’ and ‘type 2’ (DC1 and DC2, respectively) were initially described in the bone marrow (Schlitzer et al., 2015) and are now identified in all tissues in humans and mice (Guilliams et al., 2016). In general terms, DC1s are characterised by the expression of XCR1 and predominantly interact with CD8 T cells to cross-present cell-associated antigens and initiate immune responses against intracellular pathogens. DC2s, on the other hand, interact with CD4 T cells to promote T helper (Th) and T regulatory cell responses (Dudziak et al., 2007; Hilligan and Ronchese, 2020). Whilst these broad DC functions are relevant to, and conserved in, all areas of the body, the specific requirements of each tissue may also demand that these properties are appropriately modulated to promote homeostasis and optimal tissue function.

DC2s share several common characteristics that differentiate them from DC1s, including the expression of CD172a/Sirpα and being mostly depleted from the lymph nodes (LNs) of full or conditional IRF4 knockout (KO) mice (Bajana et al., 2016; Persson et al., 2013; Schlitzer et al., 2013) and humans (Guerin et al., 2018), thus validating their classification into different developmental lineages (Guilliams et al., 2014). Nonetheless, tissue DC2s are also considerably heterogeneous in phenotype, a feature that is especially notable in the migratory DC2 population that migrates from non-lymphoid tissues to secondary lymphoid organs (Alcantara-Hernandez et al., 2017; Heidkamp et al., 2016). Migratory DC2s from different tissues express unique markers that are largely conserved in mice and humans (Guilliams et al., 2016), suggesting that such heterogeneity may be acquired in response to the diverse homeostatic and functional requirements of each tissue (Sichien et al., 2017). Evidence that specific tissue-derived signals can influence the phenotype and function of local DC2s has been reported for the small intestine (SI), where retinoic acid and TGFβ have been identified as unique factors driving the development and functional maturation of intestinal CD103^+^ DC2s (Bain et al., 2017; Zeng et al., 2016). It remains unclear whether and which tissue-derived signals guide DC2 development at other sites.

Unlike the intestinal tract and airway, which are specialized for nutrient absorption and gas exchange and as such must tolerate the resulting exposure to many antigenic stimuli, the skin’s function is to block exchange thereby reducing water and nutrient loss to maintain the body’s internal environment. The skin’s barrier function is collectively carried out by several layers of tightly packed epithelial cells and sebaceous gland secretions forming an impermeable barrier. In addition, local immune cells and neural circuits perceive external threats via sensing of tissue damage, IgE-dependent mast cell activation and IL-31-dependent itch responses to control biting insects, remove ectoparasites, and limit contact with plant irritants (Bagci and Ruzicka, 2018; Olomski et al., 2020; Perner et al., 2020; Prussin and Metcalfe, 2003). Concomitantly, the skin’s continued exposure to pathogens and commensals must be managed to ensure that local invasion is prevented, tissue is repaired, and barrier function is quickly restored (Harrison et al., 2019). It is plausible that these specialized skin functions are supported by unique skin networks that are distinct from those in tissues such as lung and gut (Ronchese et al., 2020).

The skin harbours four subsets of migratory dendritic and dendritic-like cells each expressing distinct transcriptomic signatures (Miller et al., 2012). Langerhans cells represent a unique epidermal DC-like population which is developmentally distinct from conventional DCs and specialises in tolerance induction (Shklovskaya et al., 2011). Dermal DCs include DC1 and DC2 populations, with the latter being further divided into CD11b^hi^ and CD11b^low^-expressing DC2s (Henri et al., 2010). DC2s represent the most abundant dermal DC population in both mice and humans (Henri et al., 2010; Segura et al., 2012). They are mostly found in the vicinity of hair follicles but also in the deeper layer of the skin, where they form specialised niches with other immune cells including innate lymphoid cells (ILCs) (Dahlgren et al., 2019). Dermal CD11b^hi^ DC2s have been shown to induce T regulatory and anti-microbial immune responses (Naik et al., 2015; Tordesillas et al., 2018), whereas CD11b^low^ DC2s are highly responsive to Thymic Stromal Lymphopoietin (TSLP) and have been associated with driving type 2 immunity and allergic responses (Ochiai et al., 2014; Tussiwand et al., 2015). As the CD11b^low^ DC2 phenotype is unique to skin, their study may provide insight into the signals shaping the skin immune environment.

Here we report that the development of CD11b^low^ dermal DC2s requires canonical signalling through the type II IL-4 and IL-13 receptor and signal transducer and activator of transcription 6 (STAT6), and is associated with the steady-state expression of Interleukin-13 (IL-13) by ICOS^+^ KLRG1^-^ ST2^-^ ILC2s in naïve skin. No similar steady-state IL-13 expression and signaling in DC2s were observed in naïve lung or SI. In humans, IL-13-regulated genes were preferentially enriched in DC2s from healthy skin, but not lung or blood, suggesting a cytokine environment similar to murine skin. IL-13 signaling in dermal DCs promoted optimal IL-4 responses to a helminth allergen but was inhibitory to IL-17A responses to a fungal pathogen. Thus, our work identifies a skin-specific, IL-13-mediated signalling circuit between an ILC subset and DC2 precursors which is required for the development of the distinctive CD11b^low^ skin DC2 subset and establishes an anti-inflammatory, pro-Th2 environment in skin.

## RESULTS

### The transcriptomic signature of CD11b^low^ migratory DC2s in skin-draining LN (dLN) is enriched in STAT6-binding sites

To identify factors controlling the tissue-specific programming of DC2s, we developed a flow cytometric staining panel to define distinct migratory DC2 subsets in the dLNs of the skin, lung and SI. Monocytes, macrophages and neutrophils were excluded through their respective expression of Ly6C, CD64 and Ly6G, migratory DCs were identified by their expression of intermediate CD11c and high MHCII, and DC1s, Langerhans cells and DC2s were distinguished by their respective expression of XCR1, CD326 and Sirpα (**Figure S1A**) (Guilliams et al., 2016). High-dimensional analysis of concatenated samples confirmed that DC2s from different LNs expressed several unique surface markers with limited overlap and could be defined as CD11b^hi^ and CD11b^low^ DC2s in the skin-dLN, CD24+ and CD24-DC2s in the lung-dLN, and CD103+ and CD103-DC2s in the SI-dLN (**Figures 1A** and **1B**). These same phenotypes were also observed in the respective tissues of origin (**Figures 1C** and **S1B**), were present in male and female mice although in significantly different frequencies in skin (**Figure 1D**), and were observed in both BALB/c and C57BL/6 (C57) mice (**Figure 1E**), indicating that these surface markers defined conserved tissue-specific DC2 subsets under steady-state conditions. To confirm that all of these subsets were genuine bona-fide DC2s, we assessed *Irf4*^f/f^ CD11c-Cre mice in which DC2 numbers in each of the tissue-dLNs are very low (Bajana et al., 2016; Mayer et al., 2017; Persson et al., 2013; Schlitzer et al., 2013). As expected, DC2 subsets in the skin, lung and SI-dLN of *Irf4*^f/f^ CD11c-Cre mice were all significantly reduced compared to *Irf4*^f/f^ mice (**Figure S1C**).

**Figure 1.**
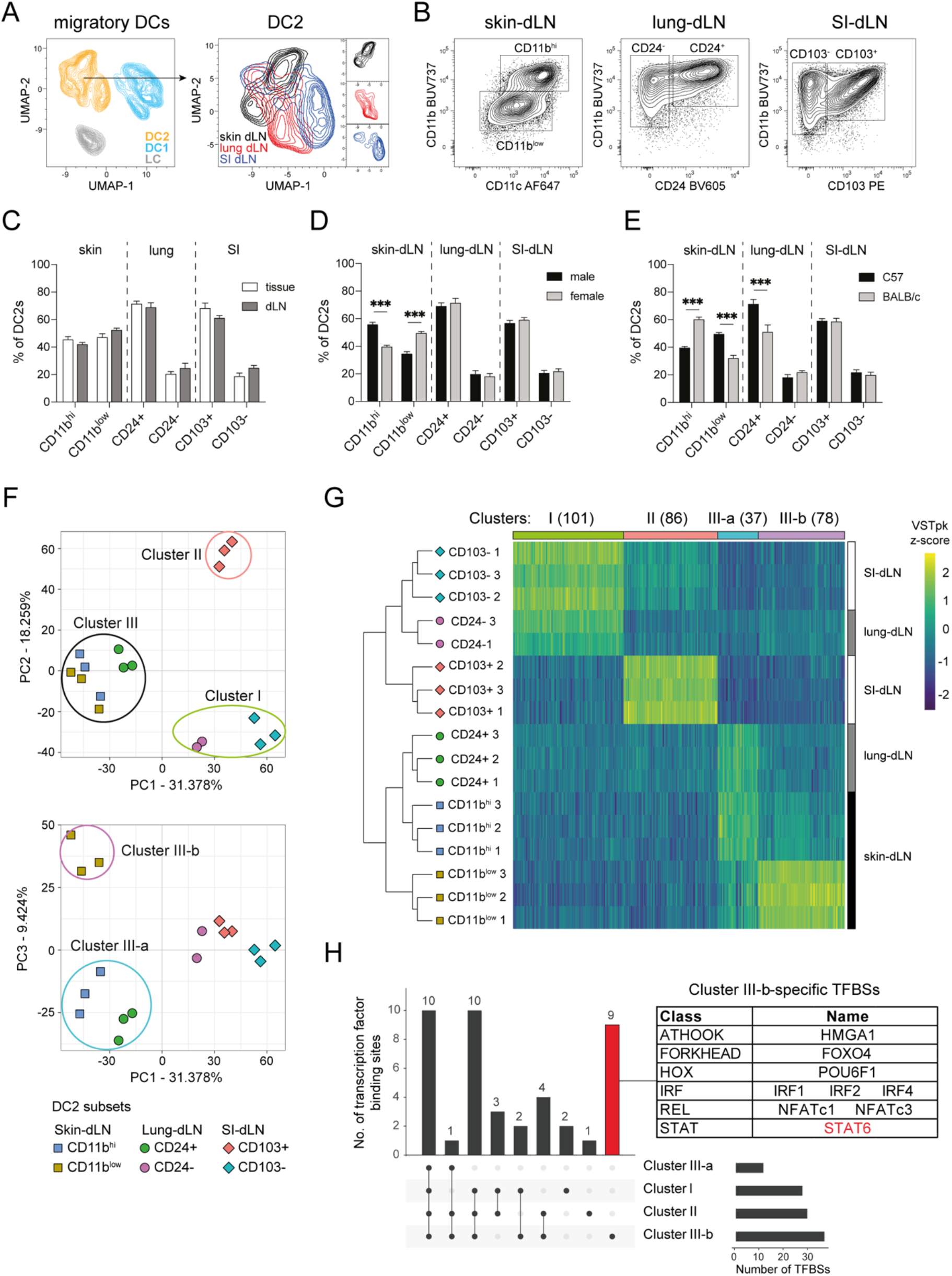
DC2s display tissue and subset-specific heterogeneity at steady state. **(A)** UMAP visualization of concatenated migratory dendritic cells (DCs) and DC2s from the skin, lung and small intestine (SI)-draining lymph nodes (dLN) of naïve C57BL/6 (C57) mice. DCs were gated as shown in Figure S1. UMAP was performed on 150,000 events (10,000 events/tissue/mouse, 5 mice per group) using 7 parameters (XCR1, CD326, Sirpα, CD11c, CD11b, CD24 and CD103) with default FlowJo settings of nearest neighbors =15 and minimum distance = 0.5 using the Euclidean distance function. **(B)** Representative contour plots displaying DC2 subsets in the skin, lung and SI-dLN. **(C)** Relative proportions of DC2 subsets within skin, lung and SI of naïve C57 mice compared to their respective dLNs. Bar graphs show mean ± SEM; each bar refers to 8-10 mice pooled from 2-3 independent experiments. **(D)** Relative proportion of DC2 subsets in the skin, lung and SI-dLN of naïve male and female C57 mice. Bar graphs show mean ± SEM; each bar refers to 6-8 mice pooled from 2 independent experiments. **(E)** Relative proportion of DC2 subsets in the skin, lung and SI-dLN of naïve C57 and BALB/cByJ (BALB/c) female mice. Bar graphs show mean ± SEM; each bar refers to 6-8 mice pooled from 2 independent experiments. (**C-E**) *P* values were determined using a two-tailed Student’s t-test. ***p < 0.001; only significant comparisons are indicated. **(F)** Principal component (PC) analysis of all expressed genes in sorted DC2 subsets from the skin, lung and SI-dLNs of naïve C57 mice reveals three main clusters in PC1 vs. PC2 and two subclusters in PC1 vs. PC3. **(G)** Heatmap of differentially expressed genes for each of the clusters and subclusters identified in (F). The number of genes in each cluster and subcluster is shown in parentheses at the top. Inclusion criteria were: protein coding genes; mean normalized RNA count > 20; fold-change greater than 2 and *p* < 0.05 as compared to each of the other clusters. VSTpk: Variance-stabilized transformed count per kbp; z-Scores for each gene were calculated using R. **(H)** UpSet plot showing the numbers of transcription factor binding sites (TFBSs) that were identified by TRANSFAC promoter analysis in each of the gene clusters in (G). The numbers of TFBSs shared across several clusters (left) or unique to individual clusters (right) are shown. The number of TFBSs that are uniquely enriched in Cluster III-b-specific genes are indicated by the red bar with the TFBSs listed in the adjacent table.

To compare the gene expression profile of each tissue-specific DC2 subset, we performed bulk RNA-sequencing of sorted migratory DC2s from the skin, lung and SI-dLN of C57 mice. A Principal Component Analysis (PCA) of all expressed genes revealed three main clusters on PC1 vs. PC2 (**Figure 1F**). Cluster I included SI-dLN CD103-DC2s and CD24-DC2s from the lung-dLN, cluster II included SI-dLN CD103+ DC2, and cluster III included both skin-dLN DC2 subsets as well as CD24+ DC2 from the lung-dLN. When PC3 was taken into account, cluster III could be further resolved into a heterogeneous cluster of skin-dLN CD11b^hi^ DC2s and lung-dLN CD24+ DC2s (cluster III-a) and a separate cluster of skin-dLN CD11b^low^ DC2s (cluster III-b). Using DESeq2 we defined a gene signature for each cluster, which included protein-coding genes that were expressed at least 2-fold higher in that cluster compared to each of the other clusters with a p-value <0.05. Based on this analysis, 101 genes were associated with cluster I, 86 with cluster II, 37 with cluster III-a, and 78 with cluster III-b (**Figure 1G** and **Table S1**).

To determine whether differential gene expression in these different DC2 clusters might be controlled by unique transcription factors, we examined the promoter sequences of the cluster-specific signature genes for enriched transcription factor binding sites (TFBS). The majority of the TFBS were shared between multiple clusters, while 2 were uniquely enriched in cluster I, 1 in cluster II and 9 in cluster III-b (**Figure 1H** and **Table S2**). No TFBS were unique to cluster III-b, possibly suggesting that these DCs may represent a ‘default’ developmental pathway. Of the 9 TFBS enriched in cluster III-b, two were for transcription factors expressed at low levels in DC2 (FOXO4, POU6F1), six were linked to interferon responses, DC2 development or T cell priming (IRF1, IRF2, IRF4, NFATc1, NFATc3, HMGA1), and one was for STAT6, a transcription factor that has no described role in the context of DC2 biology at steady state. Given that STAT6 is essential for signaling by the Th2 cytokines IL-4 and IL-13 (Ansel et al., 2006), the enrichment of STAT6-binding sites in CD11b^low^ DC2 signature genes suggested that these DC had been uniquely exposed to IL-4 and/or IL-13 in vivo.

### The development of dermal CD11b^low^ DC2s is impaired in STAT6-KO mice

To determine whether STAT6 signalling played a role in DC2 development by affecting DC2 precursors, we first identified the Lin^-^ CD11c^+^ MHCII^-^ CD135^+^ preDC population in the bone marrow (BM) and ear skin of C57 and STAT6-KO mice (**Figure S2A**). Within preDCs, the Ly6C^+^ SiglecH^-^ preDC2 population (Schlitzer et al., 2015) was present at steady state in both mouse strains in similar frequencies (**Figures 2A** and **S2A**), suggesting that STAT6 signalling did not affect preDC2 development or egress from the BM. In addition, skin preDC2s expressed similarly high levels of CD11b (**Figure 2A**), indicating that the downregulation of CD11b in dermal CD11b^low^ DC2s was occurring during maturation in the skin.

**Figure 2.**
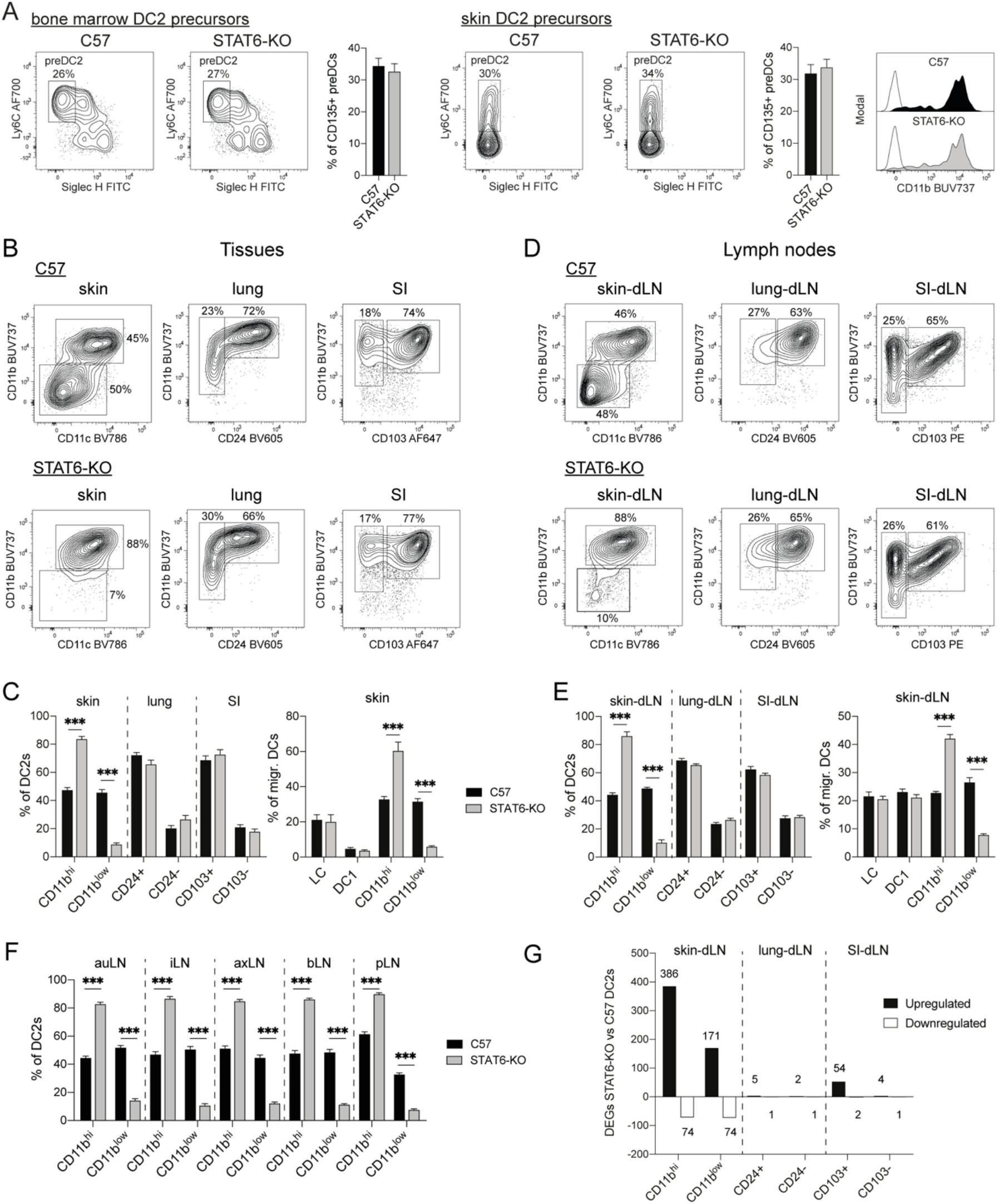
STAT6-dependent signalling controls DC2 development in the skin at steady state. **(A)** Representative contour plots showing the phenotype and frequency of preDC populations in the bone marrow and skin of naïve C57 and STAT6-KO mice, as gated in Figure S2A; preDC2 are identified by the gates and are Ly6C+ Siglec H^low^ B220^low^. Histograms on the right show CD11b expression on skin preDC2s from C57 and STAT6-KO mice (filled histograms) compared to FMO (empty histograms). Representative contour plots and histograms show concatenated data from 4-5 mice. Bar graphs show mean ± SEM; each bar refers to 6 mice (BM) or 9-10 mice (skin) pooled from 2 independent experiments. *P*=0.62 (BM) and 0.61 (ear) were determined using a two-tailed Student’s t-test. **(B)** Representative contour plots showing the phenotype and relative frequencies of DC2 subsets in the skin, lung and small intestine (SI) of naïve C57 and STAT6-KO mice. **(C)** Relative frequencies of DC2 subsets in the skin, lung and SI (left) and frequencies of DC subsets in the skin (right) of naïve C57 and STAT6-KO mice. Bar graphs show mean ± SEM, each bar refers to 7-11 mice pooled from 2-3 independent experiments. **(D)** Representative contour plots showing the phenotype of DC2 subsets in the skin, lung and SI-draining lymph nodes (dLNs) of naïve C57 and STAT6-KO mice. **(E)** Relative frequencies of migratory DC2 subsets in the skin, lung and SI-dLNs (left) and frequencies of migratory DC subsets in the skin-dLNs (right) of naïve C57 and STAT6-KO mice. Bar graphs show mean ± SEM, each bar refers to 4-5 mice (DC2 graph) or 9-10 mice (DC graph) pooled from 2 independent experiments. **(F)** Relative frequencies of CD11b^hi^ and CD11b^low^ DC2 subsets in the auricular (auLN), inguinal (iLN), axillary (axLN), brachial (bLN) and popliteal (pLN) skin-dLNs of naïve C57 and STAT6-KO mice. Bar graphs show mean ± SEM, each bar refers to 5-8 mice pooled from 2 independent experiments. (**C, E, F**) *P* values were determined using a two-tailed Student’s t-test. ***p < 0.001; only significant comparisons are indicated. **(G)** Number of up- or down-regulated differentially expressed genes (DEGs) in sorted DC2 subsets from skin, lung and SI-dLNs of naïve STAT6-KO vs C57 mice. Protein-coding genes with a mean normalized count > 20, a fold-change greater than 2 in either direction and a *p* value < 0.05 were considered differentially expressed.

Analysis of mature DC2 subsets in skin, lung and SI revealed that STAT6-KO mice exhibited a significant reduction in CD11b^low^ DC2s and a reciprocal increase in CD11b^hi^ DC2s in ear skin, while the distribution of CD24 positive and negative DC2s in the lung and CD103 positive and negative DC2s in the SI were unchanged (**Figure 2B** and **2C**). Within the skin, defective STAT6 signalling did not affect the frequency of DC1s, Langerhans cells (**Figure 2C**) or other myeloid cell populations (**Figure S2B**), pointing towards a unique requirement for STAT6 in CD11b^low^ DC2 development. Migratory DCs in the dLNs were similarly affected, with CD11b^low^ DC2s in the skin-dLN significantly reduced in STAT6-KO mice compared to C57, whereas DC2 subsets in lung or SI-dLNs, and DC1 and Langerhans cells within the skin-dLN, were present in comparable proportions (**Figures 2D** and **2E**). The findings in ear skin-dLN also applied to LNs draining other areas of the skin (**Figure 2F**), indicating that CD11b^low^ DC2s follow a similar developmental program irrespective of skin region.

To explore the impact of STAT6-dependent signaling on DC2s in further depth, we carried out a transcriptomic characterization of DC2 subsets from the skin, lung and SI-dLNs of C57 and STAT6-KO mice. Analysis of bulk RNAseq data revealed a number of differences between skin-dLN DC2 populations in STAT6-KO and C57 mice. 171 genes were upregulated and 74 downregulated in CD11b^low^ STAT6-KO DC2s compared to C57 (**Figure 2G** and **Table S3**); amongst the downregulated transcripts were *Ccl17* and *Ccl5*, two chemokines involved in functional interactions between DCs and T cells (Griffith et al., 2014), whereas transcripts for the DC lineage markers *Cd24a* and *Esam* were upregulated. In CD11b^hi^ DC2s, 386 genes were upregulated and 74 downregulated (**Figure 2G**), with many of the upregulated genes involved in transcription, motility and immune system processes. Transcripts for *Aldh1a2* which encodes Retinaldehyde Dehydrogenase 2, or RALDH2, were notably downregulated, while expression of *Ccl17* was not significantly changed in this subset. Only about 10% of the differentially expressed genes in CD11b^hi^ DC2s were similarly regulated in the CD11b^low^ population (**Figure S2C**). In line with these observations, a PCA comparing all expressed genes in skin-dLN DC2s from C57 and STAT6-KO mice showed that the separation between the two DC2 subsets was maintained across PC1, suggesting that STAT6-KO CD11b^hi^ and CD11b^low^ DC2s both maintained their transcriptional identity (**Figure S2D**), with the separation between C57 and STAT6-KO DC2s becoming apparent along PC2 for the CD11b^hi^ population and PC3 for the CD11b^lo^. In contrast to skin-dLN DC2s, lung- and SI-dLN DC2s were not strongly affected by STAT6 deficiency. In the lung-dLN, only 6 and 3 genes were differentially expressed in CD24+ and CD24-DC2s from STAT6-KO vs. C57 mice, respectively. In the SI-dLN, there were 56 genes differentially expressed in CD103+ DC2s, mostly with high *P* values, and 5 genes in CD103-DC2s (**Figure 2G**). PCAs comparing lung- or SI-dLN DC2s from C57 mice to their STAT6-KO counterparts also confirmed the overall similarity of these DC2 subsets in STAT6-KO vs. C57 mice (**Figure S2D**). Thus, STAT6 controls expression of a large number of genes in skin-derived CD11b^hi^ and CD11b^low^ DC2s under steady-state conditions.

Together, these data show that STAT6 signalling is required for the optimal development of dermal CD11b^low^ DC2 in skin, while preDC2s and DC2 subsets in lung and SI and their respective dLNs are not affected.

### ILC-derived IL-13 is necessary and sufficient for CD11b^low^ DC2 development *in vivo*

To establish whether STAT6 signalling is intrinsically required for CD11b^low^ DC2 development, we generated mixed wild-type and STAT6-KO BM chimeras in STAT6-sufficient hosts. Analysis of the skin and skin-dLNs revealed that wild-type-derived DC2s developed into CD11b^hi^ and CD11b^low^ DC2s, whereas STAT6-KO-derived DC2s were mostly CD11b^hi^ (**Figures 3A** and **S3A**), illustrating the essential intrinsic role of STAT6 for CD11b^low^ development.

**Figure 3.**
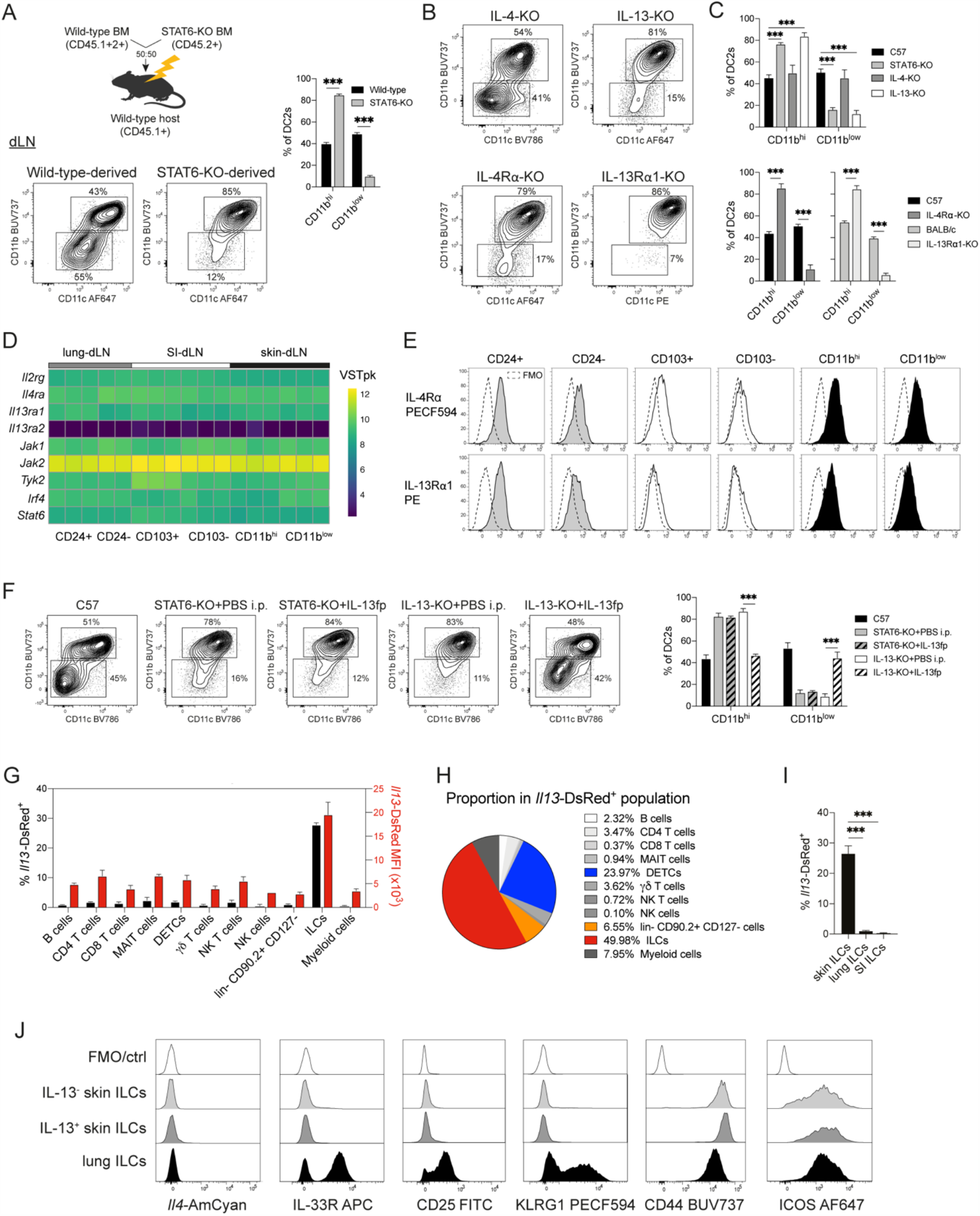
IL-13 is necessary and sufficient for the development of dermal CD11b^low^ DC2s and is expressed by skin innate cells at steady state. **(A)** Experimental set-up of mixed wild-type and STAT6-KO bone marrow (BM) chimeras. Representative contour plots show the phenotype and relative frequencies of skin-draining lymph node (dLN) DC2s in each donor BM. The bar graph shows mean ± SEM, each bar refers to 8 mice pooled from 2 independent experiments. *P* values were determined using a two-tailed Student’s t-test. ***p < 0.001. **(B)** Representative contour plots showing the phenotype and relative frequencies of skin-dLN DC2 subsets in naïve mice of the indicated strain. All KO strains were on a C57BL/6 (C57) background except for the IL-13Rα1-KO which was on a BALB/c background. **(C)** Relative frequencies of skin-dLN DC2s in naïve mice of the indicated strains. Bar graphs show mean ± SEM, each bar refers to 6-10 mice pooled from 2 independent experiments. *P* values were determined using two-way ANOVA with Sidak correction. ***p < 0.001; only significant comparisons are indicated. **(D)** Heatmap showing Variance-stabilized and transformed read counts/kbp (VSTpk) for transcripts associated with the IL-4 and IL-13 signaling pathway as assessed in sorted DC2 subsets from the lung, skin and SI-dLNs of C57 mice. **(E)** IL-4Rα and IL-13Rα1 expression on DC2 subsets from the lung, skin and SI-dLNs of C57 mice as determined by fluorescent staining and flow cytometry. Dashed lines show FMOs. Histograms show concatenated data from 4 mice in one of two independent experiments that gave similar results. **(F)** Representative contour plots showing the phenotype and relative frequencies of skin-dLN DC2s in C57 controls and in mutant mice of the indicated strains that were treated with either PBS or IL-13 fusion protein (IL-13fp) for 4 days before analysis. The bar graph shows mean ± SEM, each bar refers to 3-6 mice pooled from 1-2 independent experiments. *P* values were determined using two-way ANOVA with Sidak correction. ***p < 0.001; only relevant significant comparisons are indicated. **(G)** Frequencies of *Il13*-DsRed^+^ cells and *Il13*-DsRed Median Fluorescence Intensities (MFI) within different populations of innate and adaptive immune cells in the ear skin of naïve 4C13R reporter mice. Bar graph shows mean ± SEM, each bar refers to 6 pools of 3 mice from 2 independent experiments. *P* values were determined using one-way ANOVA with Sidak correction. ***p < 0.001; only significant comparisons are indicated. **(H)** Pie chart showing the proportion of different immune cell populations within the *Il13-*DsRed^+^ population in the ear skin of naïve 4C13R reporter mice. Data are from (G). **(I)** Frequencies of *Il13-*DsRed^+^ cells within the innate lymphoid cell (ILC) population in skin, lung and SI of naïve 4C13R reporter mice. Bar graph shows mean ± SEM, each bar refers to 6-7 mice pooled from 2 independent experiments. *P* values were determined using an ordinary one-way ANOVA with Sidak correction. ***p < 0.001; only significant comparisons are indicated. **(J)** Expression of *Il4-*AmCyan and ILC surface markers in *Il13-*DsRed-negative or -positive ILCs from the skin or lung of naïve 4C13R reporter mice. A non-reporter mice was used as *Il4-*AmCyan fluorescence-minus-one (FMO) control. Histograms show concatenated data from 4 mice in one of 4 independent experiments that gave similar results.

The canonical activation of STAT6 is mediated by the cytokines IL-4 or IL-13 that respectively signal through the IL-4Rα/common-*γ* (*γ*c) chain type I IL-4 receptor or the IL-4Rα/IL-13Rα1 type II IL-4 and IL-13 receptor (Wills-Karp and Finkelman, 2008), although alternative modes of STAT6 activation have also been reported (Chen et al., 2011). To address whether IL-4 or IL-13 signalling were responsible for STAT6 activation in dermal DC2s, we assessed IL-4-KO, IL-13-KO (**Figure S3B**), IL-4Rα-KO and IL-13Rα1-KO mice. We observed that IL-4-KO mice displayed a normal distribution of CD11b^hi^ and CD11b^low^ DC2s in the skin and minimal alterations in skin-dLN, whereas IL-13-KO mice demonstrated a pronounced reduction in CD11b^low^ DC2s that was comparable to the level observed in STAT6-KO mice (**Figures 3B, C** and **S3C, D**). CD11b^low^ DC2s were also greatly reduced in the skin and skin-dLN of IL-4Rα-KO and IL-13Rα1-KO mice, indicating that IL-13 signalling via the IL-4Rα/IL-13Rα1 type II receptor leads to the activation of STAT6 and is essential for the full development of CD11b^low^ DC2s in the skin. Transcripts for *Il2rg, Il4ra, Il13ra1* and other members of the IL-13 signalling pathway (McCormick and Heller, 2015) were expressed at similar levels by all DC2 subsets from the lung, SI and skin-dLN (**Figure 3D**), and IL-4Rα and IL-13Rα1 protein expression was low but detectable by flow cytometry at varying levels in all DC2 populations (**Figure 3E**). However, defects in STAT6 signalling only affected DC2 development in the skin and skin-dLNs, suggesting that local IL-13 production rather than differential receptor expression was driving the development of CD11b^low^ DC2s at steady state. To establish whether this was the case, we treated STAT6-KO and IL-13-KO mice with recombinant IL-13 fusion protein (IL-13fp) over 4 days. Injection of IL-13fp did not affect the proportion of CD11b^low^ and CD11b^hi^ DC2s in STAT6-KO mice, but rescued the development of CD11b^low^ DC2s in IL-13-KO mice to proportions similar to those in C57 mice (**Figure 3F**). Thus, IL-13 is both necessary and sufficient to drive the development of dermal CD11b^low^ DC2s via the activation of STAT6 under steady-state conditions.

Previous reports have shown that skin ILC2s produce low amounts of IL-13 at steady state (Roediger et al., 2013), although no biological function for such IL-13 was identified. We undertook a thorough characterization of adaptive and innate cell populations in the skin of 4C13R mice that report *Il4* expression through AmCyan and *Il13* expression through DsRed (**Figure S3E, F**). We found that almost 30% of skin ILCs were *Il13*-DsRed^+^, while other innate and adaptive immune cells were less than 5% *Il13-*DsRed^+^ (**Figure 3G**). Skin ILCs also expressed the highest level of *Il13*-DsRed per cell compared to other popoulations, and were the most abundant *Il13-*DsRed^+^ cell population at steady state followed by Dendritic Epidermal T cells (DETCs) (**Figure 3H**). Consistent with the possibility of ILCs being the source of steady-state IL-13, RAG1-KO mice showed no impairment in the development of CD11b^hi^ and CD11b^low^ DC2s in the skin and skin-dLN (**Figure S3G**), indicating that adaptive and innate-like immune cells including CD4+ T cells, DETCs, MAIT cells and NK-T cells were not essential for the steady-state production of IL-13.

Steady-state *Il13-*DsRed^+^ ILCs were identified only in the skin and not in the lung or SI (**Figure 3I**), which is consistent with our observation that the transcriptome of DC2s in lung and SI was not affected by STAT6 inactivation. *Il13-*DsRed^+^ and *Il13-*DsRed^-^ ILCs in skin and lungs did not express *Il4*-AmCyan and were CD44^hi^ and ICOS^+^. Compared to lung ILCs, skin ILCs prepared and acquired under the same conditions expressed low levels of the IL-33 receptor ST2, low CD25 (**Figure 3J**) and were mostly KLRG1- (**Figure S3F**). This observation is consistent with the described tissue adaptation and distinct expression of activating receptors on ILC2 from different tissues (Ricardo-Gonzalez et al., 2018). Together, these data are consistent with resting ILCs being the likely source of steady-state IL-13 in skin.

### The IL-13-dependent development of CD11b^low^ DC2 *in vitro* and *in vivo* requires KLF4 expression

To address whether recombinant IL-13 (rIL-13) was sufficient to drive the development of CD11b^low^ DC2s *in vitro*, we generated bone marrow derived DCs (BMDCs) in the presence of FLT3L and added rIL-13 to the culture medium at different time points and concentrations. Under standard culturing conditions, B220^-^ FLT3L BMDCs develop into XCR1^+^ DC1 and Sirpa^+^ CD11b^hi^ DC2 populations that resemble splenic DCs at steady state (Naik et al., 2007) (**Figure S4A**). We observed that increasing concentrations of rIL-13 (ranging from 1 to 50ng/ml) added on day 6 to day 9 of culture resulted in a dose-dependent development of a CD11b^low^ BMDC2 population (**Figure S4B**). A time-course experiment revealed that rIL-13 treatment was required for at least 48 hours, with the highest proportion of CD11b^low^ BMDC2s observed after the addition of rIL-13 over the last 72 hours (**Figure S4C**).

Similar to our observations *in vivo*, IL-13-dependent development of CD11b^low^ BMDC2s *in vitro* required the expression of IL-4Rα and STAT6 (**Figure 4A**). To establish whether the similarity between *in vitro* and *in vivo* DC2s extended to genes other than CD11b, we selected a small group of genes that are differentially expressed in CD11b^hi^ vs. CD11b^low^ DC2s in our RNA-seq data and assessed their expression in BMDCs. CD11b^hi^ DC2s from skin-dLN and CD11b^hi^ BMDC2s both expressed higher levels of *Itgax, Ccl9, Abcg3* and *Socs3* transcripts compared to their respective CD11b^low^ counterparts (**Figure 4B**), while *Sned1, Khdc1a, Efna5* and *Clmn* were more highly expressed in CD11b^low^ DC2s *in vivo* and *in vitro* (**Figure 4C**). Thus, rIL-13 is sufficient to drive the *in vitro* development of a CD11b^low^ DC2 population that shares transcriptional features with *in vivo* dermal CD11b^low^ DC2s.

**Figure 4.**
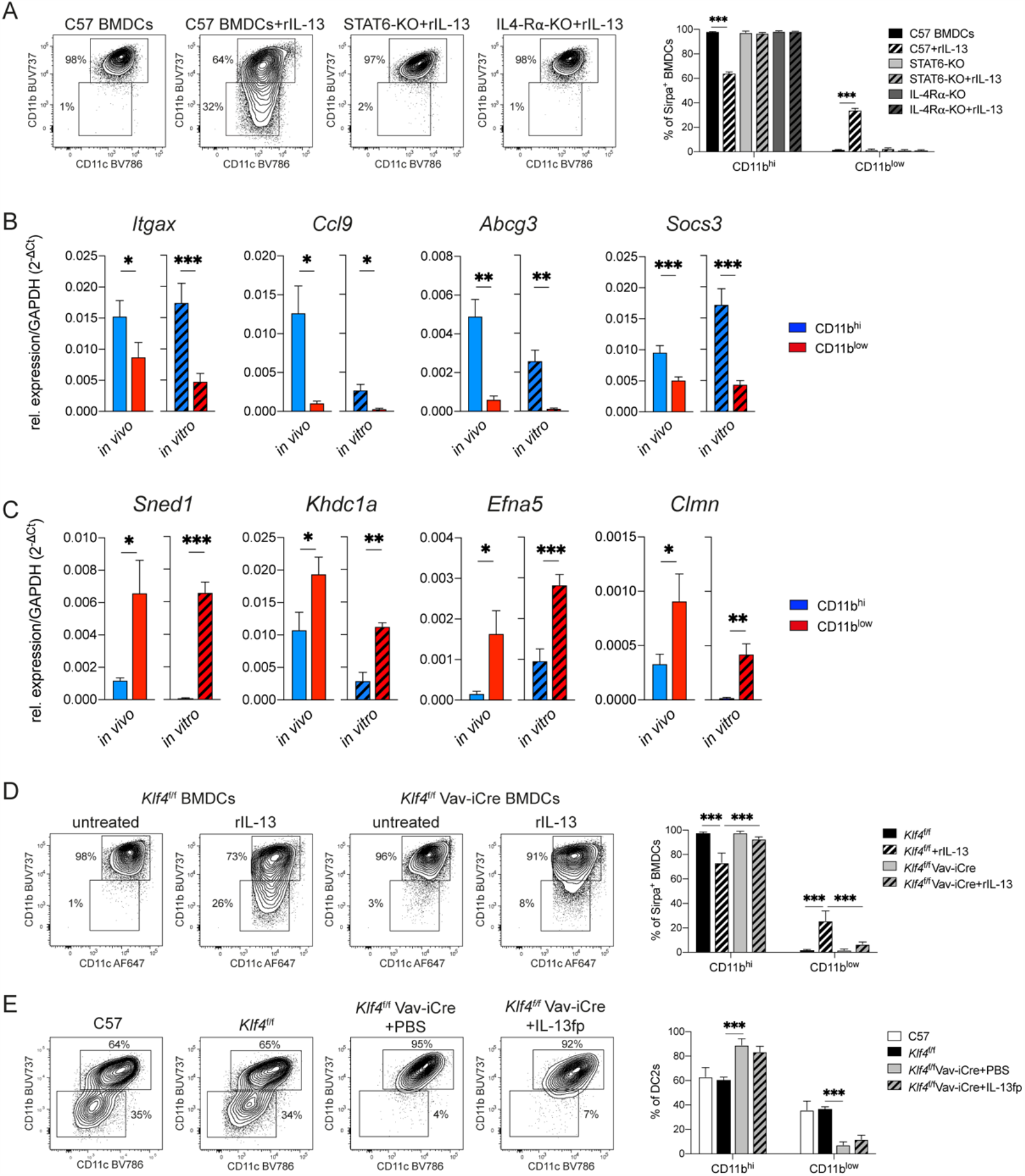
IL-13 signalling drives the development of CD11b^low^ DC2 *in vitro* and is dependent on KLF4 expression *in vitro* and *in vivo*. (**A**) Representative contour plots showing the phenotype and relative frequencies of CD11b^hi^ and CD11b^low^ Sirpa^+^ cells in FLT3L bone marrow (BM)-DC cultures from C57BL/6 (C57) mice or mice of the indicated mouse strains. Cultures were untreated or supplemented with 10 ng/ml recombinant mouse (r)IL-13 for the last 72h of culture. Bar graphs show mean ± SEM, each bar refers to 7-8 culture wells pooled from 2 independent experiments. *P* values were determined using a two-way ANOVA with Sidak correction. ***p < 0.001; only relevant significant comparisons are indicated. (**B**,**C**) Relative expression of CD11b^hi^ (B) and CD11b^low^ (C) DC2 signature transcripts in sorted DC2 populations from C57 skin-draining lymph nodes (dLN) (*in vivo*) or C57 FLT3L BMDC cultures that were supplemented with 10 ng/ml rIL-13 for the last 72h (*in vitro*). Transcript levels were measured by RT-qPCR. Bar graphs show mean ± SEM, each bar refers to 6 mice (*in vivo*) or 8 cultures (*in vitro*) pooled from 2 independent experiments. *P* values were determined using a paired, two-tailed Student’s t-test. *p < 0.05; **p < 0.01;***p < 0.001. **(D)** Representative contour plots showing phenotype and relative frequencies of CD11b^hi^ and CD11b^low^ Sirpa^+^ FLT3L BMDCs in cultures of BM cells from *Klf4*^f/f^ → C57BL/6 or *Klf4*^f/f^ Vav-iCre → C57BL/6 male chimeric mice. Cultured cells were untreated or supplemented with 10 ng/ml rIL-13 for the last 72h of culture. Bar graphs show mean ± SEM, each bar refers to 4 cultures pooled from 3 independent experiments. *P* values were determined using two-way ANOVA with Sidak correction. ***p < 0.001; only relevant significant comparisons are indicated. **(E)** Representative contour plots showing the phenotype and relative frequencies of DC2 subsets in the skin-dLNs of intact C57 mice or chimeric *Klf4*^f/f^ → C57BL/6 or *Klf4*^f/f^ Vav-iCre → C57BL/6 mice. Mice were treated with PBS or IL-13 fusion protein (IL-13fp) for 4 days as indicated. Bar graphs show mean ± SEM, each bar refers to 4-9 mice pooled from 2-3 independent experiments. *P* values were determined using two-way ANOVA with Sidak correction. ***p < 0.001; only relevant significant comparisons are indicated.

Previous reports have shown that the development of dermal CD11b^low^ DC2s requires expression of the transcription factor Krüppel-like factor 4 (KLF4) (Tussiwand et al., 2015). To establish the relationship between KLF4 and IL-13 signaling in the development of CD11b^low^ DC2s, we assessed the impact of IL-13 supplementation on FLT3L BMDC cultures from either *Klf4*^f/f^ or *Klf4*^f/f^ Vav-iCre mice. Supplementation of *Klf4*^f/f^ cultures with rIL-13 resulted in the development of a CD11b^low^ population that was similar in frequency to the population observed in BMDC cultures from C57 mice. In contrast, rIL-13 was unable to drive the development of CD11b^low^ BMDC2s in *Klf4*^f/f^ Vav-iCre BMDC cultures (**Figure 4D**), suggesting that IL-13 signaling in DC2s requires expression of KLF4. Similarly, *in vivo* IL-13fp treatment over 4 days failed to rescue the development of CD11b^low^ DC2s in *Klf4*^f/f^ Vav-iCre →C57BL/6 chimeric mice (**Figure 4E**), despite surface expression of IL-4Rα and IL-13Rα1 proteins being similar in *Klf4*^f/f^ and *Klf4*^f/f^ Vav-iCre DC2s (**Figure S4D**). As expected, *Klf4*^f/f^→C57BL/6 control chimeras displayed a normal distribution of CD11b^hi^ and CD11b^low^ DC2s in the skin-dLN. To further investigate the mechanism through which KLF4 regulates STAT6-dependent signaling, we carried out phospho-immunofluorescence experiments to compare STAT6 Y641 phosphorylation in *Klf4*^f/f^ and *Klf4*^f/f^ Vav-iCre FLT3L BMDCs exposed to rIL-13. Phosphorylated STAT6 was readily detectable in the nuclear areas of C57, *Klf4*^f/f^ and *Klf4*^f/f^ Vav-iCre Sirpa+ BMDCs 30 min after exposure to IL-13, but was not detected in IL-4Rα-KO, STAT6-KO or untreated BMDC cultures (**Figure S4E** and **S4F**), indicating that the phosphorylation of STAT6 is functional in KLF4-deficient DC2s.

Together, these results indicate that KLF4 controls DC2 responsiveness to IL-13 *in vitro* and *in vivo*, providing a mechanism for the comparable dermal DC2 phenotype observed in mice when the KLF4 and IL-13 signaling pathways are inactivated.

### IL-13 signaling in DC2s is required for optimal development of IL-4+ T cells after skin immunization

Having identified that IL-13-mediated signalling is required for CD11b^low^ DC2 development in the skin, we established two models to investigate the impact of this signalling on dermal T cell immune responses. In the first model we generated mixed BM chimeric mice to examine the functional interaction of DC and T cell populations originating from two different donor BMs (**Figure 5A**). In these chimeras, CD4^+^ T cells originated from CD45.1+2+ MHCII-KO BM and were STAT6-sufficient. In contrast, MHCII-competent DCs originated from CD45.2^+^ BMs and were either STAT6-KO and unable to respond to IL-13 signaling in the test chimeras, or WT and responsive to IL-13 in the control chimeras (**Figures 5B** and **S5A-C**). All chimeras were immunized intradermally with inactivated pathogens that induced either Th1, Th2 or Th17 responses (Blecher-Gonen et al., 2019; Hilligan et al., 2020) and CD4^+^ T cell cytokine production was assessed in the skin-dLN five days later. IFN*γ*^+^ responses induced by *Mycobacterium smegmatis* (*Ms*) immunization were detectable at similar levels in both C57 and STAT6-KO mixed BM chimeras (**Figure 5C**). In contrast, IL-4^+^ and IL-13^+^ Th cells were significantly reduced in STAT6-KO mixed BM chimeras immunized with *Nippostrongylus brasiliensis* L3 larvae (*Nb*) (**Figure 5C** and **S5C**). IL-17A^+^ Th cell responses after *Candida albicans* (*Ca*) immunization were trending to an increase in the STAT6-KO mixed BM chimeras, indicating that IL-13 signaling in DCs is important for the induction of Th2 cells but may be limiting the differentiation of Th17 cells (**Figure 5C**).

**Figure 5.**
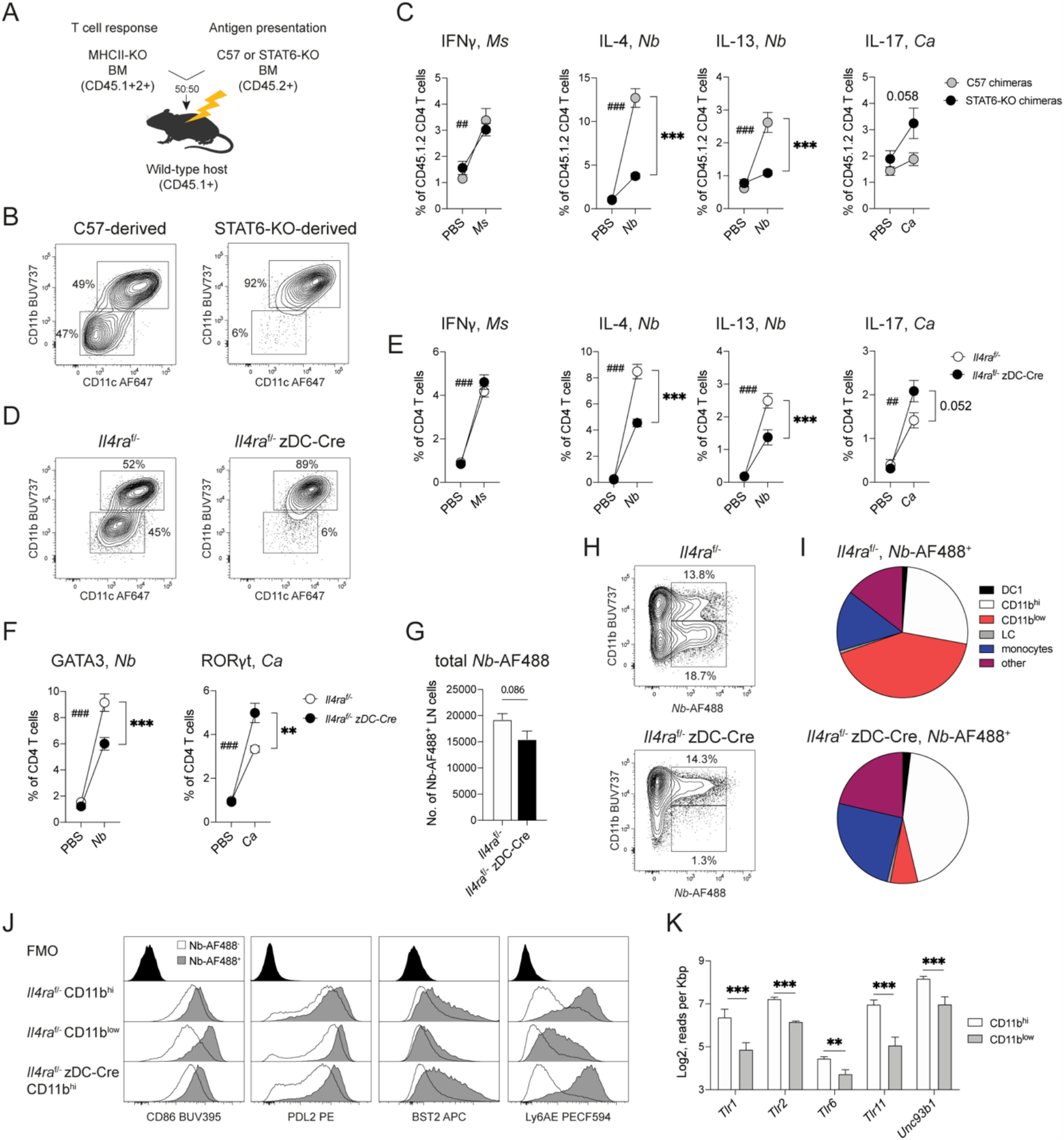
IL-13 signalling in DCs is required for optimal IL-4^+^ and IL-13^+^ CD4 T cell responses in skin-draining lymph node (dLN). **(A)** Experimental set-up of mixed bone marrow (BM) chimeras in which the antigen-presenting function of MHCII^+^ CD45.2^+^ conventional DCs from either C57 or STAT6-KO BM was assessed by measuring the response of STAT6-sufficient CD45.1^+^2^+^ T cells from MHCII-KO BM. **(B)** Representative contour plots showing phenotype and relative frequencies of C57- or STAT6-KO-derived DC2 subset in the skin-dLN of naïve C57 or STAT6-KO BM chimeras. **(C)** Frequency of cytokine-expressing CD45.1^+^2^+^ CD4 T cells in skin-dLNs of C57 or STAT6-KO mixed BM chimeras 5 days after intradermal immunization with either *Mycobacterium smegmatis* (*Ms*), *Nippostrongylus brasiliensis* L3 larvae (*Nb*), *Candida albicans* (*Ca*), or PBS as a control. Intracelluar cytokine staining was performed after 5h PMA+ionomycin stimulation. Symbols show mean ± SEM and refer to 4-6 mice/group pooled from 2 independent experiments. *P* values were determined using two-way ANOVA with Sidak correction. **(D)** Representative contour plots showing phenotype and relative frequencies of DC2 subsets in the skin-dLN of naïve *Il4ra*^f/-^ or zDC-Cre *Il4ra*^f/-^ mice. **(E)** Frequency of cytokine-expressing CD4 T cells in the skin-dLN cells of *Il4ra*^f/-^ or *Il4ra*^f/-^ zDC-Cre mice 5 days after intradermal immunization with *Ms, Nb, Ca* or PBS as a control. Intracelluar cytokine staining was performed after 5h PMA+ionomycin stimulation. Symbols show mean ± SEM and refer to 8-11 mice/group pooled from 2-3 independent experiments. *P* values were determined using a two-way ANOVA with Sidak correction. **(F)** Frequency of GATA3 and ROR*γ*t-expressing CD4 T cells in the skin-dLN cells of *Il4ra*^f/-^ or *Il4ra*^f/-^ zDC-Cre male mice 5 days after intradermal immunization with *Nb, Ca*, or PBS as a control. Intracelluar transcription factor staining was without PMA+ionomycin stimulation. Symbols show mean ± SEM and refer to 6-8 mice/group from 2 independent experiments. *P* values were determined using two-way ANOVA with Sidak correction. (**C**,**E**,**F**) * refer to the comparisons between C57 and STAT6-KO chimeras, # refer to the comparison between PBS and immunized. *p<0.05; **^,##^p < 0.01; ***^,###^p < 0.001; only significant comparisons are indicated. **(G)** Total number of AF488^+^ cells in the skin-dLN of *Il4ra*^f/-^ or *Il4ra*^f/-^ zDC-Cre mice 48 h after intradermal immunization with AF488-labelled *Nb* (*Nb*-AF488). Bar graph shows mean ± SEM, each bar refers to 7-9 mice pooled from 2 independent experiments. *P* value was determined using a two-tailed Student’s t-test. **(H)** Representative contour plots showing *Nb*-AF488^+^ uptake by DC2 subsets in the skin-dLN of *Il4ra*^f/-^ or *Il4ra*^f/-^ zDC-Cre mice 48 hours after intradermal immunization with *Nb*-AF488. **(I)** Pie charts showing the cellular composition of the AF488^+^ population in the skin-dLN of *Il4ra*^f/-^ or *Il4ra*^f/-^ zDC-Cre mice 48 hours after intradermal immunization with *Nb*-AF488. Data refer to the experiments in (G). **(J)** Histograms showing the expression of costimulatory molecules and Interferon-induced surface markers on *Nb*-AF488^-^ and *Nb*-AF488^+^ DC2 populations from *Il4ra*^f/-^ or *Il4ra*^f/-^ zDC-Cre mice 48 hours after intradermal immunization with *Nb*-AF488. Data are concatenated from 3 mice in one of two independent experiments that gave similar results. **(K)** Expression of selected *Tlr* and *Unc93b1* transcripts in DC2 subsets from the skin-dLN of naive C57BL/6 mice as determined by RNA sequencing. Bar graph shows mean ± SD, each bar refers to 3 samples each comprising the pooled LNs of 3 mice. *P* values were calculated by DESeq2; **p < 0.01; ***p < 0.001.

To confirm these results in a non-chimeric *in vivo* mouse model, we generated *Il4ra*^f/-^ zDC-Cre mice in which expression of the IL-4 and IL-13 receptors is specifically deleted in DCs thereby impairing the development of CD11b^low^ DC2s in skin and skin-dLNs (**Figure 5D**).

Upon immunization with *Ms*, we observed similar IFN*γ*^+^ CD4^+^ T cell responses in *Il4ra*^f/-^ zDC-Cre mice compared to *Il4ra*^f/-^, while IL-4^+^ and IL-13^+^ responses to *Nb* were reduced and IL-17A^+^ responses against *Ca* were trending to an increase (**Figures 5E** and **S5D**). Expression of the transcription factors GATA3 and ROR*γ*t in *Nb* or *Ca* immunized mice, respectively, reproduced the pattern observed with cytokines (**Figure 5F**). The pronounced reduction of Th2 responses upon selective depletion of IL-4 and IL-13 signalling in DC2s confirmed our observations in STAT6-KO mixed chimeric mice, and recapitulated the findings in *Klf4*^f/f^ CD11c-Cre mice (Tussiwand et al., 2015) in which impaired development of CD11b^low^ DC2s resulted in reduced type 2 immunity after skin immunization.

To assess whether the suboptimal induction of Th2 responses in *Il4ra*^f/-^ zDC-Cre mice was due to changes in DC2 number or phenotype, we injected *Il4ra*^f/-^ zDC-Cre and *Il4ra*^f/-^ mice with fluorescently-labelled *Nb* (*Nb*-AF488) to identify antigen-positive cells in dLN. The total number of AF488^+^ cells was only moderately decreased in *Il4ra*^f/-^ zDC-Cre mice compared to *Il4ra*^f/-^ (**Figure 5G**), but the composition of the AF488^+^ DC2 populations (**Figure 5H**) and the total AF488^+^ populations (**Figure 5I**) differed between the two strains. The majority of AF488^+^ cells in *Il4ra*^f/-^ mice were CD11b^low^ DC2s followed by CD11b^hi^ DC2s and monocytes. Compared to *Il4ra*^f/-^ mice, AF488^+^ cells in *Il4ra*^f/-^ zDC-Cre mice comprised a lower number of DC2s which were mainly CD11b^hi^, and an increased number of monocytes (**Figure S5E-G**). As monocyte depletion does not affect IL-4+ Th response to *Nb* (Hilligan et al., 2020), this increased monocyte response is unlikely to contribute to the Th response in *Il4ra*^f/-^ zDC-Cre mice.

The DC signals that drive the development of Th2 responses remain undefined (Lamiable et al., 2020). We therefore examined the expression of several functional activation markers on DCs (Connor et al., 2014; Connor et al., 2017; Harris et al., 1999; Pellefigues et al., 2017). In *Il4ra*^f/-^ mice, expression of the costimulatory molecules CD86 and PDL2 was higher on AF488^+^ compared to AF488^-^ DC2s, and on CD11b^low^ compared to CD11b^hi^ DC2s (**Figure 5J**), a pattern of expression similar to that observed in C57 mice (Connor et al., 2017). In *Il4ra*^f/-^ zDC-Cre mice, AF488^+^ CD11b^hi^ DC2s displayed a similar surface marker expression profile as their *Il4ra*^f/-^ counterparts, while the number of AF488^+^ CD11b^low^ DC2s was very low and insufficient for analysis. Expression of the Interferon response genes BST2 and Ly6A/E on CD11b^hi^ DC2s was also comparable in *Il4ra*^f/-^ zDC-Cre and *Il4ra*^f/-^ mice (**Figure 5J**). These results suggest that conditional inactivation of IL-4 and IL-13 signaling does not impair the response of CD11b^hi^ DC2s to *Nb* immunization, but prevents the contribution of the highly stimulatory CD11b^lo^ subset.

To better understand the increased responses to *Ca* in *Il4ra*^f/-^ zDC-Cre mice, we compared expression of relevant transcripts in the CD11b^hi^ and CD11b^lo^ DC2 subsets. As shown in **Figure 5K**, expression of the Toll-like receptor (TLR) transcripts *Tlr1, Tlr2, Tlr6* and *Tlr11* and the TLR chaperone *Unc93b1* was higher in CD11b^hi^ compared to CD11b^lo^ DC2s. As these TLRs are involved in the recognition of bacterial and fungal components and the production of pro-inflammatory cytokines including IL-6 and IL-12p40 upon recognition of fungal ligands (Pietrella et al., 2006), a higher representation of the CD11b^hi^ subset, as in the*Il4ra*^f/-^ zDC-Cre mice, could favour IL-17A responses in these mice.

### The transcriptome of human skin DC2s is enriched in IL-4/IL-13 signature genes

Although classical DC2s have been identified in humans (Dutertre et al., 2019), a human equivalent of the CD11b^low^ subset has not been described. To assess whether skin-specific IL-4 and IL-13 signalling pathways could be observed also in human DC2s, we identifed four published transcriptomic datasets that include tissue DC2s from the skin or lung of healthy individuals and compared their transcriptomic signature to the signature of control DC2s from blood or spleen within the same study (GSE35457, GSE85305 and (Chen et al., 2020) skin; GSE43184 lung).

Gene Set Enrichment Analysis (GSEA) (Subramanian et al., 2005) and the Reactome database (Jassal et al., 2020) allow for the unbiased identification of manually curated pathways that are preferentially enriched in datasets of interest. By comparing human tissue DC2s to their respective blood or spleen controls, we found that the majority of enriched pathways were unique to each individual comparison, reflecting inter-study variations (**Figure 6A**). Only one pathway, the IL-10 signaling pathway, was enriched in all four comparisons. In addition, four pathways were significantly enriched only in the three comparisons of human skin DC2s (or total DCs) to blood or spleen DC2s (or DCs); these included the ‘IL-4 and IL-13 signalling’, ‘Signaling by PDGF’, ‘TRIF TICAM1 mediated TLR4 signaling’ and ‘Regulation of insulin like growth factor’ pathways. GSEA enrichment plots for ‘IL-4 and IL-13 signaling’ genes further indicated that skin, but not lung, DC2s showed a significant enrichment for this pathway (**Figure 6B**). Heatmaps showing expression of the GSEA core-enrichment genes from the ‘IL-4 and IL-13 signalling’ pathway across the three microarray and one scRNA-seq studies revealed that similar transcripts were enriched in skin DC2 datasets compared to blood, spleen or lung DC2s (**Figure 6C** and **Figure S6A**). Interestingly, *IRF4* was enriched in human skin DC2s compared to blood, spleen or lung DC2s, and was also enriched in murine skin-dLN CD11b^low^ DC2s compared to other DC2s (**Figure 3D**), suggesting further similarities between human and murine DC2s in the skin.

**Figure 6.**
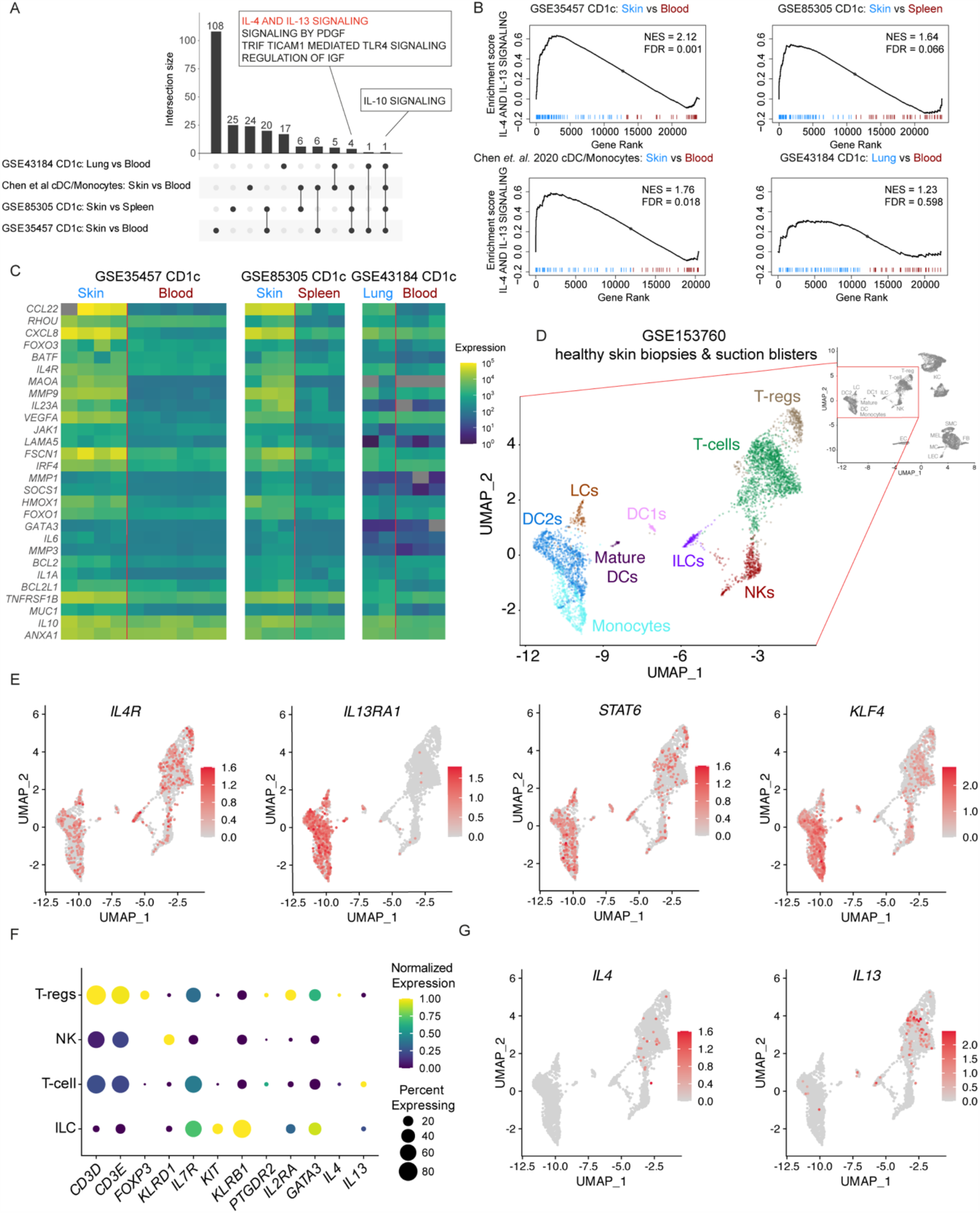
The transcriptome of DC2s from human skin, but not lung, is enriched in IL-4/IL-13 signature genes. **(A)** UpSet plot showing the numbers of Reactome pathways that are enriched in the transcriptome of sorted CD1c+ DC2s from lung vs blood (GSE43184), cDC/monocytes from from skin vs blood (Chen et al, 2020), CD1c+ DC2s from skin vs spleen (GSE85305) and CD1c+ DC2s from skin vs blood (GSE35457) as determined using Gene Set Enrichment Analysis (GSEA). All samples were from healthy donors. Pathways that are enriched only in skin samples (4 pathways) or in all comparisons (1 pathway) are listed. **(B)** GSEA enrichment plots of IL-4 and IL-13 signaling pathway genes in human tissue DC2s compared to blood or spleen as described in (A). **(C)** Heatmaps showing expression of GSEA core-enrichment genes of the IL-4 and IL-13 Reactome pathway in sorted CD1c+ DC2 from healthy human skin or lung compared to blood or spleen in the indicated microarray studies. **(D)** Detail of a UMAP plot showing lymphocyte and myeloid cell clusters from scRNA-seq of skin biopsies and suction blisters of healthy controls from the published dataset GSE153760. DC1: conventional DC1s; DC2: conventional DC2s, ILC: Innate lymphoid cells; LCs: Langerhans cells; NK: Natural killer cells. The full UMAP plot in gray includes all the clusters and can be found in Fig S6B. **(E)** Feature plots of the UMAP clusters in (D) showing the expression levels of *IL4R, IL13RA1, STAT6* and *KLF4* transcripts. Color intensity represents the level of normalized gene expression. **(F)** Dot plot showing the expression of discriminatory markers for the different lymphocyte populations shown in (D). **(G)** Feature plots of the UMAP clusters shown in (D) showing the expression levels of *IL4* and *IL13* transcripts. Color intensity represents the level of normalized gene expression.

Previous studies have detected expression of *IL13* transcripts in human skin from healthy indivdiuals (He et al., 2020; Rojahn et al., 2020), although it was unclear whether it was associated with T cells or ILCs due to the transcriptional similarity of these populations. We interrogated the dataset from one of these studies, which includes scRNA-seq data from skin biopsies and suction blisters of healthy individuals (Rojahn et al., 2020), for expression of IL-13 signalling genes in the immune and non-immune skin cell compartments. Similar to Rojahn et al, we identified several clusters of myeloid cells including monocytes, cDC1s, cDC2s, LCs and a small cluster of *CCR7*^high^/mature DCs (**Figure 6D** and **S6B-C**). Several genes that we identified as important for the IL-13-dependent development of DC2s in murine skin were also expressed in this myeloid cell compartment including *Il13RA1*, which was more highly expressed on myeloid cells than any other skin cell population, *IL4R, STAT6* and *KLF4* (**Figure 6E** and **S6C**). In the lymphocyte compartment, we identified four clusters corresponding to T-cells (*CD3*+ *FOXP3*-), T-regs (*CD3*+ *FOXP3*+), NK cells (*CD3*-*KLRD1*+) and ILCs (*CD3*-*KLRD1*-*IL7R*^high^ and *KLRB1*+) (**Figure 6F**). Cells within the ILC cluster also expressed *KIT, IL2RA, GATA3*, which have been associated with KIT+ ILC3s and IL2RA+ GATA3+ ILC2s (Teunissen et al., 2014). Unlike flow cytometry (Wojno et al., 2015), *PTGDR2* encoding the prostaglandin D2 receptor CRTH2 was not sufficient to identify ILC2s by scRNA-seq due to its low expression. *IL13* expression was detected at low level in 5.12% and 2.96% of the T cell and ILC clusters in healthy skin, respectively. By contrast, only T cells, but not ILCs, expressed *IL4* (**Figure 6F-G**).

Taken together, our reanalysis of five published human datasets shows that human DC2s from healthy adult skin express an IL-4/IL-13 gene signature compared to DC2s from other tissues, suggesting that the transcriptome of human skin DC2s could be influenced by local IL-13 as demonstrated in mice.

## DISCUSSION

In this paper we report that the steady-state production of IL-13 in murine dermis is necessary and sufficient for the development of a population of dermal DC2s that is unique to skin and is characterized by low expression of CD11b. We show that skin-migratory CD11b^low^ DC2s express a STAT6-dependent signature and that their development is impaired in mice that are either STAT6 deficient, IL-13 deficient or do not express a functional IL-13 receptor. The inability of DC2s to sense IL-13 in STAT6-KO or IL-4Rα conditional KO mice is functionally consequential and results in impaired IL-4+ and increased IL-17A+ Th cell responses in skin-dLN. By re-analysing published transcriptomic data, we also show that dermal DC2s in healthy human skin express an IL-13-dependent transcriptional signature and skin ILCs express IL-13, thus suggesting that DCs in healthy murine and human skin are exposed to similar IL-13 environments in the steady-state, and might exhibit similar functional properties.

The heterogeneity of the IRF4-dependent DC2 subset across different tissues is well recognised (Alcantara-Hernandez et al., 2017; Guilliams et al., 2016; Heidkamp et al., 2016) and presumably develops in response to tissue-specific signals and growth factors that remain often uncharacterised (Sichien et al., 2017). A notable example of DC2 specialization is the CD103^+^CD11b^+^ DC2 subset unique to the SI lamina propria and mesenteric LN (Bain et al., 2017). In the absence of TGFβR signalling, SI DC2s remain CD11b^+^CD103^-^ and are unable to promote the development of FoxP3^+^ T regulatory cells and mucosal Th17 populations. As cytokines of the TGFβ family are ubiquitously expressed in epithelia and are essential for the development and function of multiple tissues, they are well suited to a role in the tissue adaptation of DC populations. Unlike TGFβs, IL-13 does not have a recognized role in tissue homeostasis. The main known function of IL-13 is its role in allergic inflammation and parasite infection (Wynn, 2003). High IL-13 production in skin, as observed in atopic disease, is associated with fibrosis, decreased expression of filaggrin and filaggrin-like molecules by keratinocytes and loss of skin barrier function (McAleer and Irvine, 2013). However, low levels of IL-13 produced by DETCs at steady state were reported to enhance epidermal integrity (Dalessandri et al., 2016), and *Il13* transcripts together with low but measurable protein levels are constitutively expressed by dermal ILC2s in healthy murine skin (Roediger et al., 2013) and freshly explanted human ILC2s (Teunissen et al., 2014), raising the question of a potential homeostatic function for IL-13 (Strid et al., 2016). Here we provide evidence that homeostatic IL-13 production in skin has a clear immunological effect, firstly by supporting dermal DC2 differentiation at steady state, and secondly by regulating the ability of DC2s to induce immune responses after immunization. To our knowledge, this is the first report of a lymphocyte-derived cytokine contributing to tissue-specific DC2 development in the steady state, and the first evidence of skin as an “IL-13-conditioned” tissue in non-pathological conditions. It remains to be established how the function of other DC and non-DC poulations in skin might also be affected by IL-13 signaling.

The DC2 phenotype in STAT6-KO and IL-4Rα conditional KO mice was similar to mice that are total or conditional KO for KLF4, a transcription factor required for the development of CD11b^low^ CD24^low^ ‘Double Negative’ dermal DC2s and for the induction of Th2 responses (Tussiwand et al., 2015). We found that the similarity between STAT6-KO and KLF4-KO DC2s was not due to STAT6 deficiency affecting expression of *Klf4* transcripts in DC2s, but to the ability of KLF4 to control the response of DC2s to IL-13 signalling *in vitro* and *in vivo*, thus revealing a previously unrecognized link between KLF4 and IL-13 signalling in DCs. The basis of the defective responsiveness to IL-13 remains unclear as expression of IL-4Rα and IL-13Rα1 and phosphorylation of STAT6 were all similar in KLF4-KO and C57 DCs, but might be explained by a transcriptional cooperation between KLF4 and STAT6 as documented in M2 macrophages (Liao et al., 2011). Futher experiments are required to determine whether a cooperation between KLF4 and STAT6 controls DC2 transcriptional responses to IL-13. Additional effects of DC-specific KLF4 deletion, such as the increased DC1/DC2 ratio and decreased DC2 numbers in multiple tissues (Tussiwand et al., 2015), were not recapitulated in STAT6-KO and IL-4Rα conditional KO mice thus may be driven by broader functions of KLF4, or its early expression during DC development and prior to preDCs exiting the BM.

Apart from a moderate difference between males and females, which was presumably due to the suppressive activity of male hormones on ILC2 function (Laffont et al., 2017), the ratio of dermal CD11b^low^ to CD11b^hi^ DC2s appeared fairly constant in different laboratories and in different mice regardless of anatomical location, implying a regulated mechanism that controls the equilibrium between the CD11b^low^ and CD11b^hi^ DC2 subsets. Treatment with recombinant IL-13 was sufficient to rescue the development of CD11b^low^ DC2s in IL-13-KO mice but did not induce downregulation of CD11b on all dermal DC2s, suggesting that additional factors may influence the differentiation of CD11b^low^ DC2s such as the positioning within an appropriate niche (Dahlgren et al., 2019) or a pre-existing heterogeneity within the DC2 population (Sichien et al., 2017). The notion that IL-13 is not the only signal controlling the development of dermal CD11b^low^ DC2 is also compatible with the presence of a small population of CD11b^low^ DC2s in STAT6-KO and IL-4Rα-KO mice and the high number of differentially expressed genes in CD11b^hi^ DC2s from STAT6-KO compared to C57 mice, with the latter suggesting that the STAT6-KO CD11b^hi^ population might include incompletely differentiated CD11b^low^ DC2s.

Our data suggest that steady-state IL-13 production is exclusive to the skin. Skin-specific expression of IL-13 by ILC2s has been reported in mice and humans (Roediger et al., 2013; Salimi et al., 2013), is consistent with the unique transcriptional profile of skin ILC2s (Ricardo-Gonzalez et al., 2018), and appears to be independent of skin microbiota or signaling from innate cytokines including TSLP, IL-33 and IL-25 ((Ricardo-Gonzalez et al., 2018) and our unpublished observations) suggesting a key role of the specific tissue environment in determining ILC function. The mechanism through which ILC-derived IL-13 conditions skin DC2s at steady state also appears to differ from the mechanisms in lung, where allergen and IL-33-dependent IL-13 production by ILC2s were necessary for DC migration to the dLN (Halim et al., 2014) and for the later recruitment and activation of Th2 cells in non-lymphoid tissue (Halim et al., 2016; Halim et al., 2018; Nieuwenhuizen et al., 2018; Oliphant et al., 2014). While a STAT6 signature was detectable in CD11b^low^ DC2s in naïve mice, our previous transcriptomic analyses of DC2 subsets in PBS and *Nb*-immunized mice failed to detect an IL-4/IL-13-dependent signature in *Nb*-treated vs PBS mice (Connor et al., 2017), suggesting that DC2s were either not exposed to, or not further affected by, these cytokines after immunization. Together, these observations suggest that, compared to lung, the skin environment is intrinsically predisposed to Th2 induction, although both skin and lung can sustain Th2 priming in the appropriate conditions. Consistent with this view, IL-13 signaling enabled the increased expression of costimulatory molecules on CD11b^low^ DC2s, both at steady state and after *Nb*-AF488 uptake. The increased responsiveness of CD11b^low^ DC2s to TSLP (Ochiai et al., 2014), an epithelial cytokine with a known role in promoting Th2 responses (Ziegler, 2012) may underpin this observation. In addition, the higher expression of several *Tlr* transcripts in CD11b^hi^ compared to CD11b^low^ DC2s suggests a lower responsiveness of CD11b^low^ DC2s to microbial stimuli, which was observed in our previous study (Hilligan et al., 2020) and is a likely consequence of IL-13 signaling (Leyva-Castillo et al., 2020), providing a plausible mechanism for the increased IL-17A and ROR*γ*t+ responses in IL-4Rα conditional KO mice.

The observation of a skewed ability of skin DC2 to induce immune responses raises the question of how the low-dose, steady-state IL-13 that imparts a pro-Th2 environment in naïve skin could be advantageous for skin homeostasis given that it appears to be conserved across mice and humans. The skin environment plays an important role in the response to commensals such as *S. epidermidis* (Harrison et al., 2019; Naik et al., 2015) in which specific T cells produce IL-17A under homeostatic conditions for antimicrobial defense and also express Th2 potential for tissue repair. Within the context of a response to commensals, IL-13-dependent DC conditioning might promote equilibrium by preventing excessive IL-17A production and consequent neutrophil infiltration, thereby avoiding damaging responses to commensals while supporting expression of repair-promoting cytokines and growth factors upon tissue injury (Harrison et al., 2019).

In addition to the potentially beneficial regulation of Th17 responses in skin, our data also reveal a mechanism for the preferential priming of Th2 responses after skin exposure. In skin, Th2 responses are reported to curb entry of helminth parasites (Obata-Ninomiya et al., 2013), or promote itch and scratch responses to insect bites or ticks to favour their removal and repair the resulting damage (Allen and Wynn, 2011; Karasuyama et al., 2020). However, Th2 responses can also cause pathology and develop into damaging allergic responses. While a preferred route for allergic sensitization in humans has not been established, epidemiological studies suggest that delayed introduction of some foods to infants may increase the likelihood of allergy development by providing greater opportunity for contact through skin (Du Toit et al., 2008). Similarly, genetic variations that decrease the barrier function of skin are associated with increased incidence of allergic disease that is not limited to skin but extends to respiratory and food allergy (Irvine et al., 2011; McAleer and Irvine, 2013). Our study provides a potential mechanism for these observations by documenting an IL-13-dependent and tissue-specific conditioning of dermal DC2s that leads to Th2-biased functional properties.

In conclusion, steady-state IL-13 production by dermal ILC2s underpins the development of a CD11b^low^ DC2 subset that is unique to skin, and preferentially directs mouse CD4^+^ T cell differentiation to an IL-4^+^ Th2 phenotype while inhibiting Th17 responses. The functional consequences of this circuit in humans, in which an IL-13-dependent signature in skin DC2 is also apparent, are yet to be characterised. However, genome-wide association studies showing an opposing impact of *IL13* gene polymorphisms in Atopic dermatitis vs Psoriasis (Baurecht et al., 2015), two skin diseases respectively dominated by dysregulated activation of Th2 and Th17 cells, respectively, suggest a similar regulation in humans and an important role of this mechanisms in the propensity to disease.

## ACKNOWLEDGEMENTS

The authors wish to thank Prof Graham Ogg, Oxford University, for sharing raw data of the scRNA-seq analysis of human skin blister (Chen et al, 2020); Dr. Patrick M. Brunner and Dr. Vera Vorstandlechner, Department of Dermatology, Medical University of Vienna, for their advice on single cell QC filtering and data analysis of scRNA-seq data of human skin biopsies and blister cells (GSE153760, Rojahn et al, 2020); Yueqi Wang, Google, for developing the Shiny browser for single cell RNAseq data, and providing debugging help, and all colleagues at the Malaghan Institute of Medical Research for discussion and suggestions. We also thank Dr. Olivier Gasser and the NIH Tetramer Facility for providing the 5-OP-RU-loaded tetramers for MAIT cell identification. The MR1 tetramer technology was developed jointly by Dr. James McCluskey, Dr. Jamie Rossjohn and Dr. David Fairlie, and the material was produced by the NIH Tetramer Core Facility as permitted to be distributed by the University of Melbourne, Australia. Finaly, we gratefully acknowledge the flow cytometry support of the members of the Hugh Green Cytometry Centre and the expert animal husbandry of the Biomedical Research Unit.

This work was funded by an Independent Research Organization grant from the Health Research Council of New Zealand (HRC) to the Malaghan Institute, a Health Research Council of NZ project grant to FR, a Malaghan Institute Postdoctoral Fellowship to KLH, the Marjorie Barclay Trust, and the Intramural Research Program of the NIAID, NIH, USA.

## AUTHOR CONTRIBUTIONS

Conceptualization: JUM, OL, FR; software: SIO, DAE; formal analysis: JUM, SIO, DAE, LM-E; Investigation: JUM, OL, KLH, JSC, RGD, JY, GRW, LME, KAW, EJH, SCT, SCC, LMC; Resources: CRM, FB, AS, RT, DJ, MRH, GLG; data curation: SIO, DAE; writing the original draft: JUM, FR; reviewing & editing the original draft: JUM, OL, FR with input from all authors; visualization JUM, OL, KLH, SIO, DAE; supervision: MRH, FR; funding acquisition: AS, FR.

## MATERIALS AND METHODS

### Mice

Mice were bred and housed under specific pathogen-free conditions at the Malaghan Institute of Medical Research and provided with water and feed *ad libitum*. All mice were between 6 and 14 weeks of age and were age- and sex-matched within experiments. C57BL/6J, BALB/cByJ, B6-SJ ptprca, CD11c-Cre (B6.Cg-Tg(Itgax-cre)1-1Reiz/J), IRF4-flox (B6.129S1-Irf4^tm1Rdf^/J), STAT6-KO (B6.129S2(C)-Stat6^tm1Gru^/J) and zDC-Cre (B6.Cg-Zbtb46^tm3.1(cre)Mnz^/J) mice were from breeding pairs originally obtained from the Jackson Laboratories, USA. 4C13R reporters (Tg(Il4-AmCyan,Il13-DsRed*)1Wep (Roediger et al., 2013) and IL-4-KO (Il4^tm1.1Wep^) (Hu-Li et al., 2001) mice were from breeding pairs kindly provided by Dr William Paul, NIAID, NIH, Bethesda USA; IL4Ra-KO (Il4ra^tm1Fbb^) (Mohrs et al., 1999) and IL-4Rα-flox (Il4ra^tm2Fbb^) (Herbert et al., 2004) were from breeding pairs kindly provided by Dr Frank Brombacher, University of Capetown, ZA; MHCII-KO (H2-Aa^tm1Blt^) (Kontgen et al., 1993) were bred from pairs originally provided by Dr Horst Bluethmann, Hoffman-La Roche Ltd, Basel, Switzerland. *Itgax*-cre^+/-^ *Irf4*^fl/fl or fl/-^ (IRF4^ΔCD11c^) and *Irf4*^fl/fl or fl/-^ (IRF4^WT^) mice were generated by crossing *Itgax*-cre to *Irf4*^fl^ for two generations. *zBTB46*-cre^+/-^ *Il4r*^fl/-^ (IRF4^ΔzDC^) and *Il4r*^fl/-^ mice were generated by crossing *Zbtb*-cre females to *Irf4*^fl^ for two generations.

For some experiments, C57BL/6Tac and BALB/c were obtained from Taconic Biosciences (Rensselaer, NY) while B6.SJL-*Ptprc*^*a*^ *Pepc*^*b*^/BoyJ, RAG1-KO on C57BL6/J background, and IL13Ra1-KO on BALB/c background were from the NIAID contract facility at Taconic Biosciences.

Experimental protocols were approved by the Victoria University of Wellington Animal Ethics Committee or the NIAID Animal Care and Use Committee and performed according to institutional guidelines.

### Generation of IL-13 KO mice

IL-13-KO mice were generated by the Australian Phenomics Network and Monash Genome Modification Platform, Melbourne, Australia, using the strategy illustrated in Figure S3B. CAS9 enzyme, guide RNA (gRNA) 1 targeting intron 1 (5’ - AGAGUCUUGGAGCUGAAAGA −3’) and gRNA2 targeting intron 3 (5’-CUUAGAGCGUUACAAGUCC-3’) of the murine *Il13* gene were injected into C57BL/6 fertilized eggs to remove exon 2 and 3. Mice were screened for exon deletion by PCR using the primers P1 (5’-GAGGCTGGCATGGTGGTTTC-3’) and P2 (5’-TGGAGACCTGTGAAACGGCA-3’). Of two founder mice carrying exon 2-3 deletions, only one gave viable progeny carrying the mutation when crossed to C57BL/6. These heterozygous mice were mated to each other to generate IL-13-KO mice carrying a homozygous exon 2-3 deletion.

### BM Chimeric Mice

BM cell suspensions were prepared by flushing cut femurs and tibias with unsupplemented IMDM (Gibco) and filtering through a sterile 70 µm cell strainer (Falcon). Recipient mice aged 10-12 weeks were treated with two doses of 5.5 Grey using a Gammacell® 3000 Elan irradiator (MDS Nordion) given 3 hours apart. One day after irradiation, mice received 1-8 × 10^6^ Klf4^fl^ Vav-iCre or Klf4^fl^ BM cells (Tussiwand et al., 2015) from gender-matched donor mice as detailed in the text and Figure legends.

For the generation of allogeneic mixed-BM chimeras, BM donor mice were depleted of T cells by intraperitoneal injection of 200 µg *InVivoMAb* anti-mouse Thy-1 (Clone T24/31, BioXCell) on day −5 and −2 before BM collection. Recipient mice were injected with 100 µg anti-mouse Thy-1 i.p. one day after BM transfer. Chimeric mice were housed in individually ventilated cages and supplied with 2 mg/mL Neomycin trisulfate-supplemented drinking water for the first 3 weeks. For steady-state DC profiling and BMDC generation, chimeric mice were used at least 8 weeks after reconstitution, or immunized after 12 weeks of reconstitution for the characterization of CD4^+^ T cell responses.

### Immunizations and *in vivo* treatments

Mice were anesthetized with 100 µg/g body weight Ketamine and 3 µg/g body weight Xylazine (both Provet) by intraperitoneal injection and injected with 4×10^6^ CFU heat-killed *Ms*, 1×10^7^ heat-killed *Ca* or 300 non-viable Nb L3 larvae in 30 µl PBS intradermally in the ear pinna as previously described (Hilligan et al., 2020). PBS was injected into the ear pinna of control animals. 60 µl of PBS or 20 µg IL-13 fusion protein (Absolute Antibody) in the same volume were injected intraperitoneally on day 0, 60 µl of PBS or 10 µg IL-13 fusion protein were injected on day 2 and 3 and tissues harvested on day 4.

### Preparation of immunogens

*M. smegmatis* (*Ms*, mc2155) was grown in LB broth (Difco LB Lennox – low salt, BD) under agitation at 37°C overnight. Bacteria were washed three times in PBS containing 0.05% Tween 80 and heat-killed at 75°C for 1 hour and stored at −70°C. *N. brasiliensis* (*Nb*) infective L3 larvae were prepared as described (Camberis et al., 2013), washed three time with sterile PBS followed by three washes with antibiotic wash buffer (AWB: PBS supplemented with 100 µg/ml Gentamcycin (Sigma-Aldrich) and 500 U/ml Penicillin-Streptomycin (Gibco)). Larvae were incubated in AWB twice for 1 hour at room temperature, washed four times with sterile PBS and inactivated by three freeze-thaw cycles. *C. albicans* (*Ca*) was propagated by inoculating sterile yeast extract-peptone-dextrose broth (Difco YPD, BD) and incubating under agitation at 30°C for 72 hours. Yeasts were washed in PBS and heat-killed at 75°C for 1 hour. For fluorescent labelling of antigens, non-viable *Nb* were incubated in 0.05 M NaHCO3 buffer and 0.1 mg AF488 NHS Ester (Molecular Probes, Invitrogen) for 15 minutes and washed with 0.1M Tris buffer according to the manufacturer’s instructions.

### Tissue sampling and processing

To prepare single cell suspensions for DC assessment, auricular or inguinal LNs draining the skin, mediastinal LNs draining the lung or mesenteric LNs specifically draining the small intestine were disrupted with 18G needles and digested in IMDM containing 100 µg/mL Liberase TL and 100 µg/mL DNase I (both Sigma-Aldrich) for 30 minutes at 37°C. Cells were collected, filtered through a 70 µm cell strainer, washed with FACS buffer (PBS supplemented with 2% FCS, 2 mM EDTA and 0.01% Sodium Azide (all Gibco)) and maintained at 4°C. For skin preparations, ears were collected and split into ventral and dorsal halves, cut into small pieces and digested in 2ml HBSS (Gibco) containing 2 mg/mL Collagenase IV (Sigma-Aldrich) and 100 µg/mL DNase I (Sigma-Aldrich) for 30 minutes at 37°C and 250 rpm in a shaking incubator. The suspension was collected, dissociated using a syringe and 18G needle, filtered through a 70 µm cell strainer, washed with FACS buffer and maintained at 4°C. Lungs were collected after perfusion, cut into small pieces and digested in 1 ml IMDM (Gibco) containing 500 µg/mL Liberase TL and 500 µg/mL DNase I (both Sigma-Aldrich) for 45 minutes at 37° C and 150 rpm in a shaking incubator. Cells were collected, filtered through a 70 µm cell strainer, washed with FACS buffer and maintained at 4°C. Small intestine preparations were performed as previously described (Ferrer-Font et al., 2020). Small intestinal segments were excised, Peyer’s patches were removed and the intestines were opened longitudinally. Segments were when washed in PBS, cut into 0.5 cm pieces, collected in cold HBSS and washed twice with HBSS containing 2 mM EDTA (both Gibco) for 15 minutes at 37° C and 200 rpm in a shaking incubator to dissociate the epithelium. Segments were then digested in RPMI (Gibco) containing 10% FCS (Gibco), 1 mg/ml Collagenase VIII and 50 µg/ml DNase I (both Sigma-Aldrich) for 15-20 minutes at 37°C and 200 rpm in a shaking incubator. The suspension was then filtered through a 100 µm and 40 µm cell strainer, washed with FACS buffer and maintained at 4°C.

Skin-dLN preparations for the assessment of T cell responses were generated by pressing the LNs through a 70 µm cell strainer to obtain single cell suspensions, washed with FACS buffer and maintained at 4°C. For the assessment of intracellular cytokines in T cells, LN single cell suspensions were incubated in TCM containing 50 ng/mL PMA (Sigma-Aldrich), 1 µg/mL ionomycin (Sigma-Aldrich) and 1 µL/mL GolgiStop (BD Pharmingen) for 5 hours before proceeding with surface and intracellular flow cytometry staining.

### FLT3L bone marrow cultures

BM cells were suspended in Red Blood Cell Lysing Buffer Hybri-Max™ (Sigma-Aldrich) for 1 min at RT, washed and resuspended in tissue culture medium (TCM: IMDM supplemented with 5% fetal calf serum (FCS), 1% Penicillin-Streptomycin and 55 µM 2-Mercaptoethanol (all Gibco) supplemented with 4% FLT3L supernatant (generated from the cell line CHO flag Flk2.clone5 kindly provided by Prof Nic Nicola, WEHI). BM cells were cultured in 6-well plates (Corning) at 5×10^6^/well in 5 ml TCM, and incubated at 37°C in 95% humidity and 5% CO2 for 9 days. Cultures were fed every three days by removing 2 mL of medium and adding 2 mL of TCM containing 10% FLT3L supernatant. Recombinant murine IL-13 (Peprotech) was added at different times and concentrations as indicated in each experiment.

### Phospho-STAT6 Immunofluorescence

At day 9, unstimulated FLT3L bone marrow DCs (BMDCs) were harvested and one million cells were resuspended in 1 mL of TCM in 24 wells plates and stimulated with recombinant mouse IL-13 (Peprotech) at a concentration of 100 ng/mL for 30 min at 37°C. 1×10^5^ cells were then cytocentrifuged onto microscope glass slides (Trajan), encircled with a PAP pen (Daido Sangyo), fixed with 4% paraformaldehyde (ThermoFisher) for 10 min at room temperature, and permeabilized with 100% methanol at 4°C for 10 min, as previously described (Schmidt et al., 2019). Blocking was performed for 1 hour at room temperature using SuperBlock T20 (ThermoFisher), before incubation with primary antibody (anti-pSTAT6 Tyr641, ThermoFisher) for 1 hour at room temperature at 1:100 dilution in PBS and washed three times with PBS. Slides were then incubated with secondary antibody anti-rabbit Alexa Fluor 647 (ThermoFisher), anti-CD172a Alexa Fluor 594 (Biolegend) and DAPI (0.5µg/ml, Merck). Slides were mounted with Faramount (Agilent) and covered with a glass coverslip (22×22, Thermofisher). Images were obtained using the FV3000 confocal microscope (Olympus) with a 10x objective, using the 405, 594 and 640 nm laser lines and 3 GaAsp photomultiplier detectors. Images were acquired with a 16-bit depth, a line average of 3 using sequential scanning and a pixel dimension of 2048×2048 (1μm=1.6091pixels). Acquired images were first pre-processed with the FIJI imaging processing package (v2.0.0-rc-69/1.52i, (Schindelin et al., 2012)). Cell masks were generated using CellProfiler (v4.0.3, (Jones et al., 2008)), while .fcs files using the cell masks and channel images were generated with Histocat (v1.76, (Schapiro et al., 2017). Gating and cell marker statistics were performed using Flowjo software (v10.7, BD), and these statistics were imported into GraphPad Prism 8 (v8.4, GraphPad Software, Inc.) for statistical analysis.

### Flow cytometry and cell sorting

Single cell suspensions were incubated in FACS buffer containing anti-mouse CD16/32 (clone 2.4G2, hybridoma culture supernatant, 1:300 dilution) to block Fc receptors prior to labelling with a mix of fluorescently labeled monoclonal antibodies for 30 minutes at 4°C. Anti-CD3 (145-2C11), anti-CD4 (RM4-5), anti-CD11b (M1/70), anti-CD11c (HL3), anti-CD326 (G8.8), anti-IA/IE (M5/114.15.2), anti-NK1.1 (PK136), anti-CD45R (RA3-6B2), anti-CD86 (GL1), anti-Ly6A-E (D7), anti-CD44 (IM7), anti-CD103 (M290) and anti-NK1.1 (PK136) were from BD Biosciences. Anti-CD64 (X54-5/7.1), anti-CD8 (2.43), anti-Ly6C (HK1.4), anti-Ly6C (AL-21), anti-CD11c (N418), anti-CD26 (H194-112), anti-CD172a (P84), anti-CD45.1 (A20), anti CD45.2 (104), anti-CD45 (30-F11), anti-CD278 (C398.4A), anti-CD90.2 (30-H12), anti-CD90.2 (53-2.1), anti-CD124 (I015F8), anti-CD278 (7E.17G9), anti-CD24 (30-F1), anti-CD25 (3C7), anti-TCR”# (GL3), anti-CD49b (DX5), anti-Ly6C/Ly6G (RB6-8C5), anti-CD200R3 (Ba13), anti-FcεRIα (MAR-1), anti-CD317 (927), anti-CD117 (2B8), anti-CD19 (6D5) and anti-XCR1 (ZET) were from BioLegend. Anti-KLRG1 (2F1), anti-CD19 (ID3), anti-Siglec-F (E50-2440), anti-TCR$ (H57-597), anti-CD127 (A7R34), anti-CD273 (TY25), anti-Ly6G (1A8), anti-CD206 (MR6F3), anti-IL33R (RMST2-33) and anti-CD213a (13MOKA) were from ThermoFisher. 5-OP-RU-loaded MR1 tetramer conjugated to BV421 was kindly provided by Dr. Olivier Gasser. Dead cells were excluded from analysis using DAPI (Molecular Probes) or ZOMBIE NIR (Biolegend). For staining of intracellular cytokines, cells were stained with LIVE/DEAD™ Fixable Blue (Molecular Probes) prior to staining of cell surface molecules. Cells were then fixed and permeabilized using a BD Cytofix/Cytoperm kit (BD Pharmingen) and stained for intracellular cytokines. Anti-IL-4 (11B11) and anti-IFN*γ* (XMG1.2) were from BD Biosciences; anti-IL-13 (eBio13A) and anti-IL-17A (eBio17B7) were from ThermoFisher.

For staining of transcription factors, cells were stained with LIVE/DEAD™ Fixable Blue (ThermoFisher) prior to staining of cell surface molecules. Cells were then fixed and permeabilized using a True-Nuclear™ Transcription Factor Buffer Set (BioLegend) and stained with anti-ROR*γ*t (Q31-378) and anti-GATA3 (C50-823), both from BD Biosciences. Single-stain controls were prepared with UltraComp eBeads (ThermoFisher) following the manufacturer’s instructions, or cells, if this resulted in a brighter signal, and were used to calculate a compensation matrix. Sample acquisition was performed on a LSR Fortessa or FACSymphony (both BD Biosciences) using BD FACS DIVA software or a 3L or 5L Aurora Spectral Flow Cytometer (Cytek Biosciences) using SpectroFlo software v2.2. Final analysis and graphical output were performed using FlowJo software (v10.7, BD), OMIQ (v2020, OMIQ, Inc.). For sorting of DC subsets for quantitative RT-PCR analysis, cells were stained as described and DC2 subsets from skin dLN were sorted as indicated in Figure S1. Cells were directly sorted into RNA lysis buffer (Zymo Research) using a BD INFLUX Cell sorter (BD Biosciences) and frozen at –80°C until RNA extraction. Three biological replicates were prepared in two separate DC sorting experiments and purity of target cell population was >95%.

For sorting of DC subsets for RNA sequencing, cells were stained as described and presorted as MHCII^hi^ migratory DCs from skin, lung and SI dLNs as indicated in Figure S1 using a BD INFLUX Cell sorter (BD Biosciences). Cells were resuspended in FACS buffer and the respective DC2 subets were sorted directly into QIAzol Lysis Reagent (QIAGEN) and frozen at –80°C until RNA extraction. Three biological replicates were prepared in three separate DC sorting experiments and purity of target cell population was >98%.

### RNA isolation and sequencing

Total RNA was prepared from frozen cell pellets using the RNeasy Micro Kit (QIAGEN). RNA was quantified using a Quantus Fluorometer (Promega) and RNA integrity was checked using a Fragment Analyser (Agilent). Library preparation and RNA sequencing were contracted out to Otago Genomics, University of Otago, NZ. Samples were spiked with External RNA Controls Consortium mix (Ambion; Life Technologies) and ribosomal RNA was depleted using RiboZero (Illumina) before library preparation. Paired-end stranded RNA sequencing was performed on an Illumina HiSeq 2500 V4 sequencing system using Illumina TruSeq kits. Between 10 and 30 million read pairs were generated per sample.

### Read mapping and differential expression analysis

Paired-end raw FASTQ files were trimmed using Trimmomatic (v0.36) (Bolger et al., 2014) and aligned using STAR (v2.7.1a) (Dobin et al., 2013) to *Mus musculus* mm10 M16 (GRCm38.p5) genome downloaded from GENCODE. Aligned reads were quality-checked using MultiQC (v1.7) (Ewels et al., 2016) and counted using the R (v3.4.4) package Rsubread (v1.28.1) (Liao et al., 2013). Differentially expressed genes were identified using the R package DESeq2 (v1.28.1) (Love et al., 2014) and the output was filtered for log2 fold-change >1 in either direction and adjusted p-values < 0.05. VSTpk (variance-stabilized reads per thousand base pairs) values were generated from DESeq2 normalised counts using the DESeq2 Variance Stabilizing Transformation with default parameters, then dividing the resulting value by the length of the longest gene isoform in kb. Visualizations were made using the R packages pheatmap (v1.0.12), UpSetR (v1.4.0) (Conway et al., 2017) and Tidyverse (v1.3.0). The cluster-specific gene signatures in Figure 1 and Table S1 were created using DESeq2 and comprised of all the protein-coding genes that were differentially expressed in a certain cluster compared to each of the other clusters, with a Fold Change > 2 in either direction and adjusted p-value < 0.05. VSTpk z-scores were calculated using the R function base::scale(center = TRUE) on VSTpk values in a column-wise fashion for each gene.

### Transcription factor binding site analysis

For *in silico* prediction of transcription factor binding sites, the promoter sequences of the genes of interest (Table S1) were analysed using TRANSFAC (Matys et al., 2006). Briefly, sequences from 5000 bp to 100 bp upstream (from −5000 bp to +100 bp) of transcription start site were selected and analysed against a random set of genes of the same size with the Immune cell specific group of matrices (p < 0.01, v2020.02).

### Quantitative RT-PCR

RNA was extracted from 1000 cells using the Quick-RNA Microprep Kit (Zymo Research) following the manufacturer’s guidelines. cDNA was synthetized from the total amount of extracted RNA using the High capacity RNA-to-cDNA kit (ThermoFisher). qPCR targets were preamplified using SsoAdvanced Preamp supermix (Bio-rad) following the manufacturer’s guidelines. As specified by the manufacturer’s guidelines, preamplified cDNA was diluted 40 times and RT-qPCR was performed using SYBR Green Master Mix (ThermoFisher) and the forward (FW) and reverse (REV) primers (IDT) FW 5’-TGCTCAGGACCAGACCAATTC-3’ and REV 5’-CCTGGAAATCTCTGCAGGTGT-3’ for *Itgax*, FW 5’-CAGATCACACATGCAACAGAGAC-3’ and REV 5’-TCAATTTCAAGCCCTTGCTGTG-3’ for *Ccl9*, FW 5’-TTCATGGACCTGCACAGGATG-3’ and REV 5’-GAAGTCTGCAGGGTTGTTGTG-3’ for *Abcg3*, FW 5’-GCGAGAAGATTCCGCTGGTA-3’ and REV 5’-CCGTTGACAGTCTTCCGACA-3’ for *Socs3*, FW 5’-GCCTGCACACCTCTACATCAT-3’ and REV 5’-CTGAGCATCCGTGAGTGGTG-3’ for *Sned1*, FW 5’-GCTGTGGGTGATGGACATGAT-3’ and REV 5’-TCCCTCAGAGGATGATCCCG-3’ for *Khdc1a*, FW 5’-GCTACGCCGTCTACTGGAAC-3’ and REV 5’-AGTGAGGGCAGAAAACATCCA-3’ for *Efna5*, FW 5’-CTCTCTGGGCGAAATCTGCT-3’ and REV 5’-GAGTGCTTTCGCTATGTTGTTCA-3’ for *Clmn* and FW 5’-TGATGGGTGTGAACCACGAG-3’ and REV 5’-GCCGTATTCATTGTCATACCAGG-3’ for *Gapdh* using a QuantStudio 7 (Applied Biosystems). Transcript levels are expressed as the ratio of 2−ΔCT (Transcript of interest)/2−ΔCT (*Gapdh*).

All primer pairs were designed in house using Primer3Plus, blasted against mouse transcriptome with Primer-BLAST and pre-tested to assess that amplicons of the corrects size were generated and produced single peaks upon melt curve analysis.

### Human DC2 transcriptomic data integration

Microarray data were downloaded from GEO: human skin and blood DC2 (healthy volunteers) (Haniffa et al., 2012) were downloaded from GEO: GSE35457, microarray data from adult human skin and spleen DC2 (healthy volunteers) (McGovern et al., 2017) were downloaded from GSE85305, and microarray data from human lung and blood DC2 (healthy volunteers) (Yu et al., 2013) were downloaded from GSE43184. Microarray data were annotated using the ‘illuminaHumanv4.db’ R package (v1.26.0). Single cell RNA-seq data of human skin blister and blood cDC/Monocytes from healthy volunteers (Chen et al., 2020) were obtained from the authors and normalised using the SCTransform function of Seurat v3.2.0 (Stuart et al., 2019). In order to compare with the microarray data, gene expression for each group was summarised as the mean SCT expression across all cells within the group. Human skin blister cells and biopsy from healthy volunteers (Rojahn et al., 2020) were downloaded from GEO (GSE153760) and processed using Seurat v3.2.0 (Stuart et al., 2019), following the v3.2 “Integration and Label Transfer” vignette. Cells were pre-filtered to include only cells with at least 500 features and less than 10000 counts, as well as restricting mitochondrial fraction to between 0.5 and 15 percent. Filtered cells were then transformed within-sample via SCTransform, followed by integration using the 3,000 most variable features. UMAP dimensionality reduction was carried out, using 50 principle components and running for 400 epochs. One cluster exhibiting high variance in the UMAP plot was removed, prior to visualising the UMAP, coloured by both cluster and expression for selected genes. Feature and dot plots were created using tools integrated within Seurat.

### Gene Set Enrichment analysis

GSEA (Subramanian et al., 2005) and the Reactome database (Jassal et al., 2020) were used to identify enriched signalling pathways in human DC2 from published microarray and scRNA-seq studies. Results from GSEA were processed using a custom script to recover cumulative per-marker scores for all ranked genes, so that they could be replotted with identical scales.

### Data access

Raw RNA-seq data as FASTQ files have been deposited in NCBI SRA under bioproject PRJNA668222. All code and associated parameters are available at https://github.com/fronchese/Mayer_et_al_2020.

### Quantification and statistical analyses

Statistical analyses were performed using Prism 8.0 GraphPad software. Number of samples, independent experiments and statistical tests used are indicated in the figure legends. Data were analysed using ordinary one-way ANOVA when comparing multiple groups, two-way ANOVA with Sidak’s multiple comparisons test when comparing two variables, or a two-tailed Student’s *t*-test when comparing two groups as indicated in each figure legend. Statistical significance was defined as p < 0.05.

---

**Figure S1.**
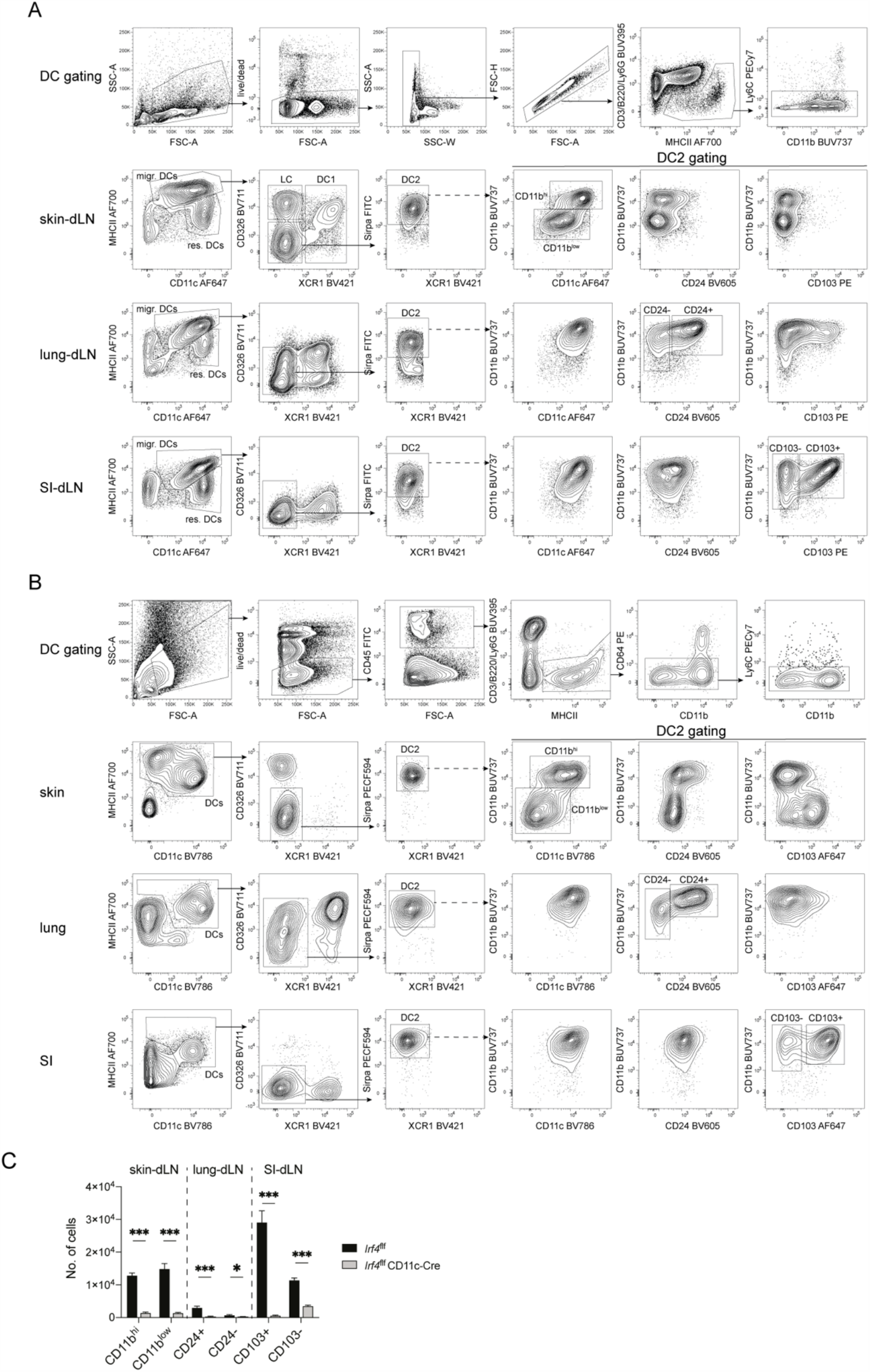
Related to Figure 1: Gating strategy for DC and DC2 populations in the skin, lung and small intestine (SI) of naïve mice and their corresponding draining lymph nodes (dLNs). **(A)** Gating strategy for migratory DC2 subsets in dLNs. Generic gating on live single CD3^-^ B220^-^ Ly6G^-^ Ly6C^-^ cells was performed for all samples before gating on dLN-specific DC2 subsets. **(B)** Gating strategy for DC2 subsets in tissues. Generic gating on live single CD45^+^ CD3^-^ B220^-^ Ly6G^-^ CD64^-^ Ly6C^-^ cells was performed for all samples before gating on tissue-specific DC2 subsets. **(C)** Numbers of DC2s in the skin, lung and SI-dLN of *Irf4*^f/f^ and *Irf4*^f/f^ CD11c-Cre mice. Bar graphs show mean ± SEM, each bar refers to 5-8 mice pooled from 2 independent experiments. *P* values were determined using a two-tailed Student’s t-test. *p < 0.05; ***p < 0.001; only significant comparisons are indicated.

**Figure S2.**
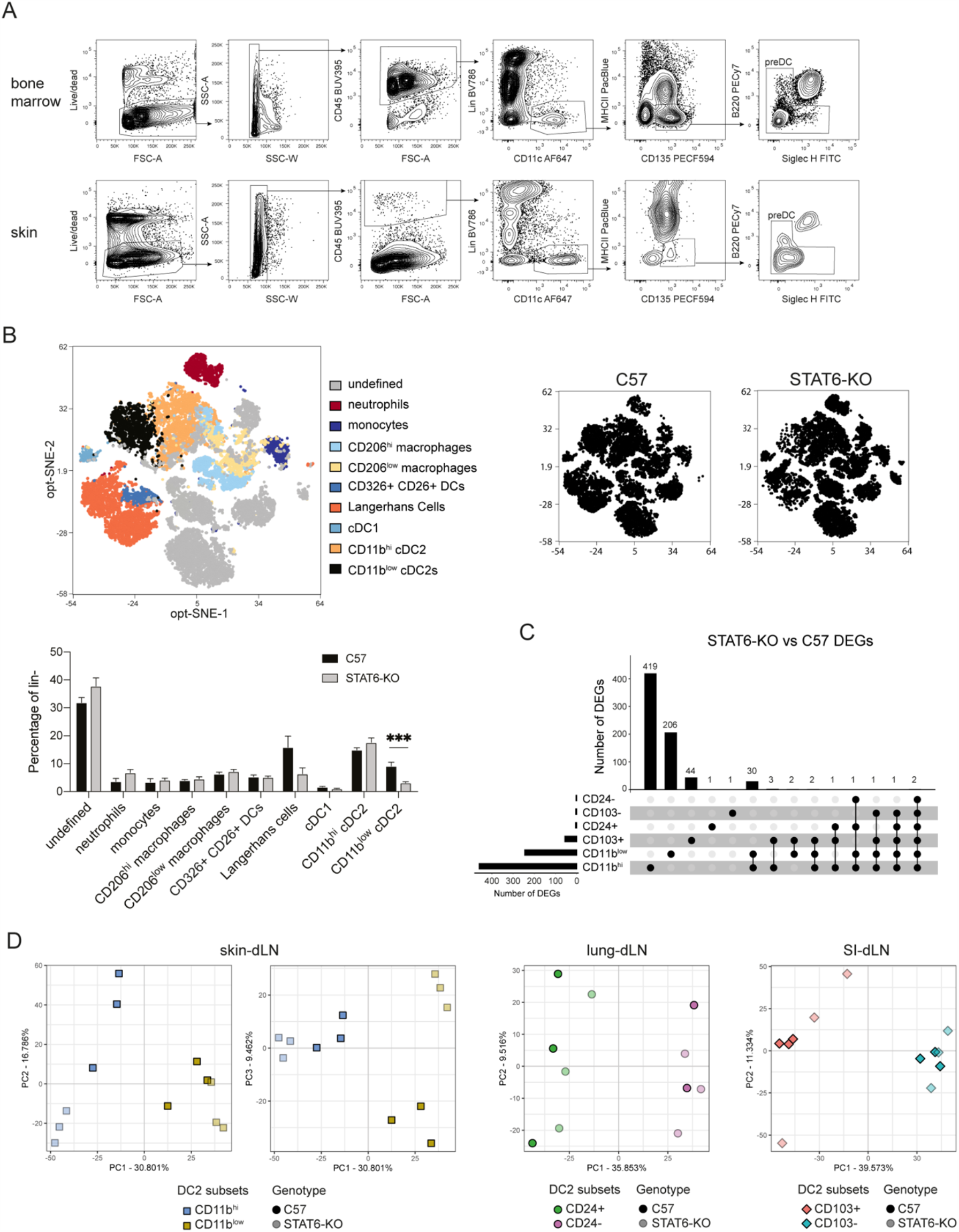
Relating to Figure 2: STAT6 signalling controls a distinct transcriptional program in CD11b^hi^ and CD11b^low^ DC2s. **(A)** Gating strategy for preDC2s in the bone marrow (upper panels) and skin (lower panels). **(B)** Representative tsne visualization of concatenated live single CD45^+^ CD3^-^ CD19^-^ myeloid cells from the skin of naïve C57BL/6 (C57) and STAT6-KO male mice; combined data from both strains are shown in the color image. Opt-SNE was performed on 16648 events (2081 events/mouse, 4 mice per genoype) using 12 parameters (XCR1, CD206, CD11b, Ly6c, CD11c, Ly6G, SIRPα, CD326, CD26, CD64, BST2, MHC-II) with default OMIQ settings and a perplexity of 30 with 1000 iterations. The bar graph shows mean ± SEM, each bar refers to 9 mice pooled from 2 independent experiments. *P* values were determined using a two-tailed Student’s t-test. ***p < 0.001; only significant comparisons are indicated. **(C)** UpSet plot showing the number of unique and shared differentially expressed genes (DEGs, including up- and down-regulated) in the indicated DC2 subsets sorted from the skin, lung and SI-draining lymph nodes (dLNs) of STAT6-KO and C57 mice using the gating in S1A. **(D)** Principal component (PC) analyses of all expressed genes in sorted DC2 subsets from the skin (upper panels), lung and SI (lower panels) −dLNs of naïve C57 and STAT6-KO mice.

**Figure S3.**
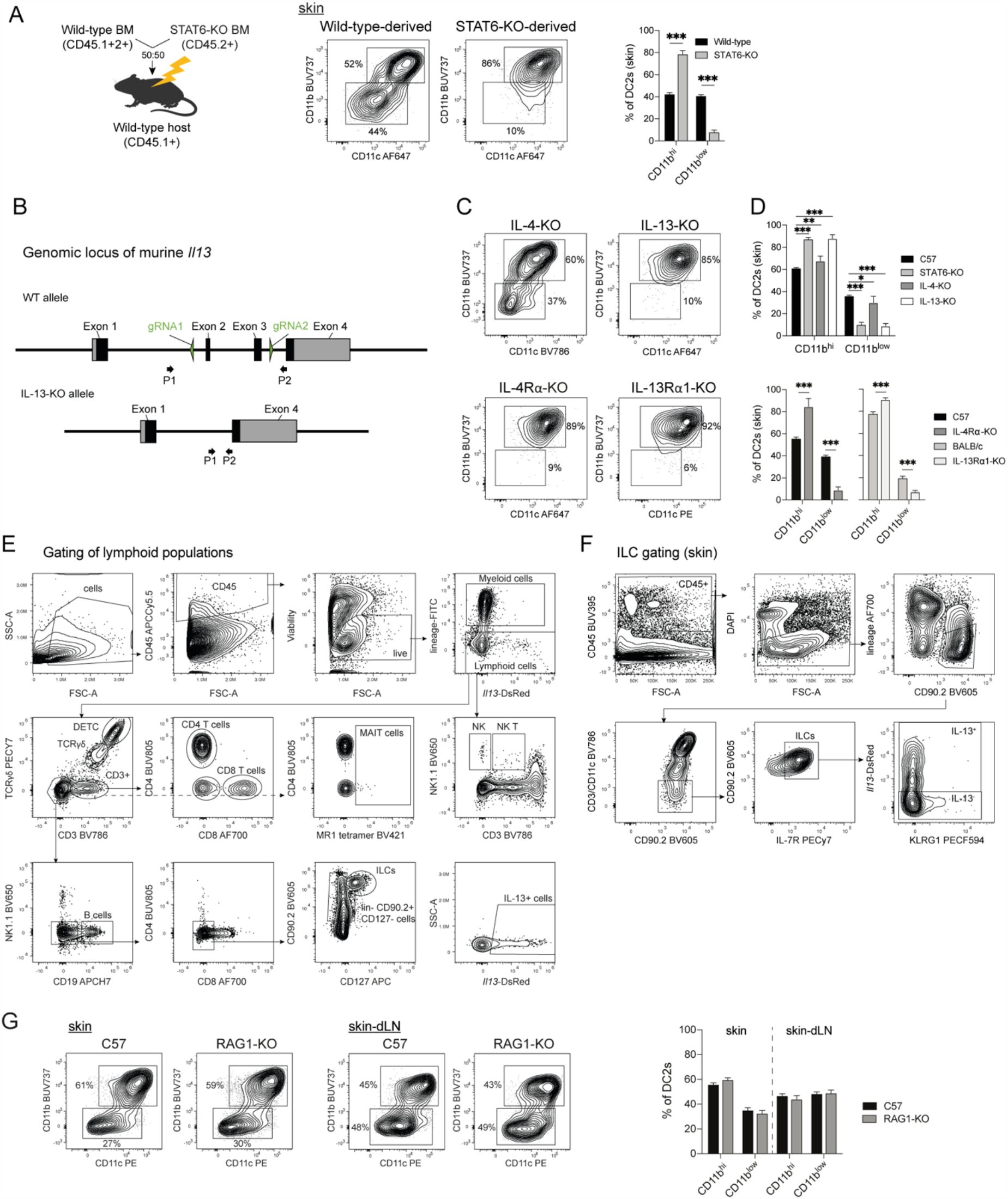
Related to Figure 3: IL-13 signalling is necessary for the development of CD11b^low^ DC2s in skin and is independent of adaptive or KLRG1+ cell popoulations. **(A)** Experimental set-up of the mixed wild-type and STAT6-KO BM chimeras in Figure 3A-C. Representative contour plots show the phenotype and relative frequencies of skin DC2s from each donor BM. The bar graph shows mean ± SEM, each bar refers to 8 mice pooled from 2 independent experiments. *P* values were determined using a two-tailed Student’s t-test. ***p < 0.001. **(B)** Schematic of the genomic *Il13* locus of IL-13-KO mice illustrating the deletion of exons 2 and 3. **(C)** Representative contour plots showing the phenotype and relative frequencies of skin DC2s in naïve mice of the indicated strains. All KO strains were on a C57BL/6 background except for the IL-13Rα1-KO which were on a BALB/c background. **(D)** Relative frequencies of skin DC2 subsets in naïve mice of the indicated strains. Bar graphs show mean ± SEM, each bar refers to 5-15 mice pooled from 2-3 independent experiments. *P* values were determined using two-way ANOVA with Sidak correction. *p < 0.05; **p < 0.01; ***p < 0.001. **(E)** Gating strategy to identify lymphoid cell populations in the ear skin of naïve 4C13R reporter mice. To define cells of lymphoid origin, cells were pre-gated on single, live, CD45^+^, myeloid-lineage^-^ (CD11b^-^ Ly6G^-^ Ly6C^-^ CD11c^-^) cells. *Il13-*DsRed^+^ cells were gated within each cell population, or in the total CD45^+^ population as shown in the lower right panel. **(F)** Gating strategy to identify innate lymphoid cells (ILCs) in the skin of naïve 4C13R reporter mice. A similar gating strategy was used to identify ILCs in lung and small intestine. **(G)** Representative contour plots showing the phenotype and relative frequencies of skin and skin-draining lymph node (dLN) DC2 subsets in naïve C57BL/6 (C57) and RAG1-KO mice. Bar graph shows mean ± SEM, each bar refers to 8-10 mice pooled from 2 independent experiments. *P* values were determined using a two-tailed Student’s t-test.

**Figure S4.**
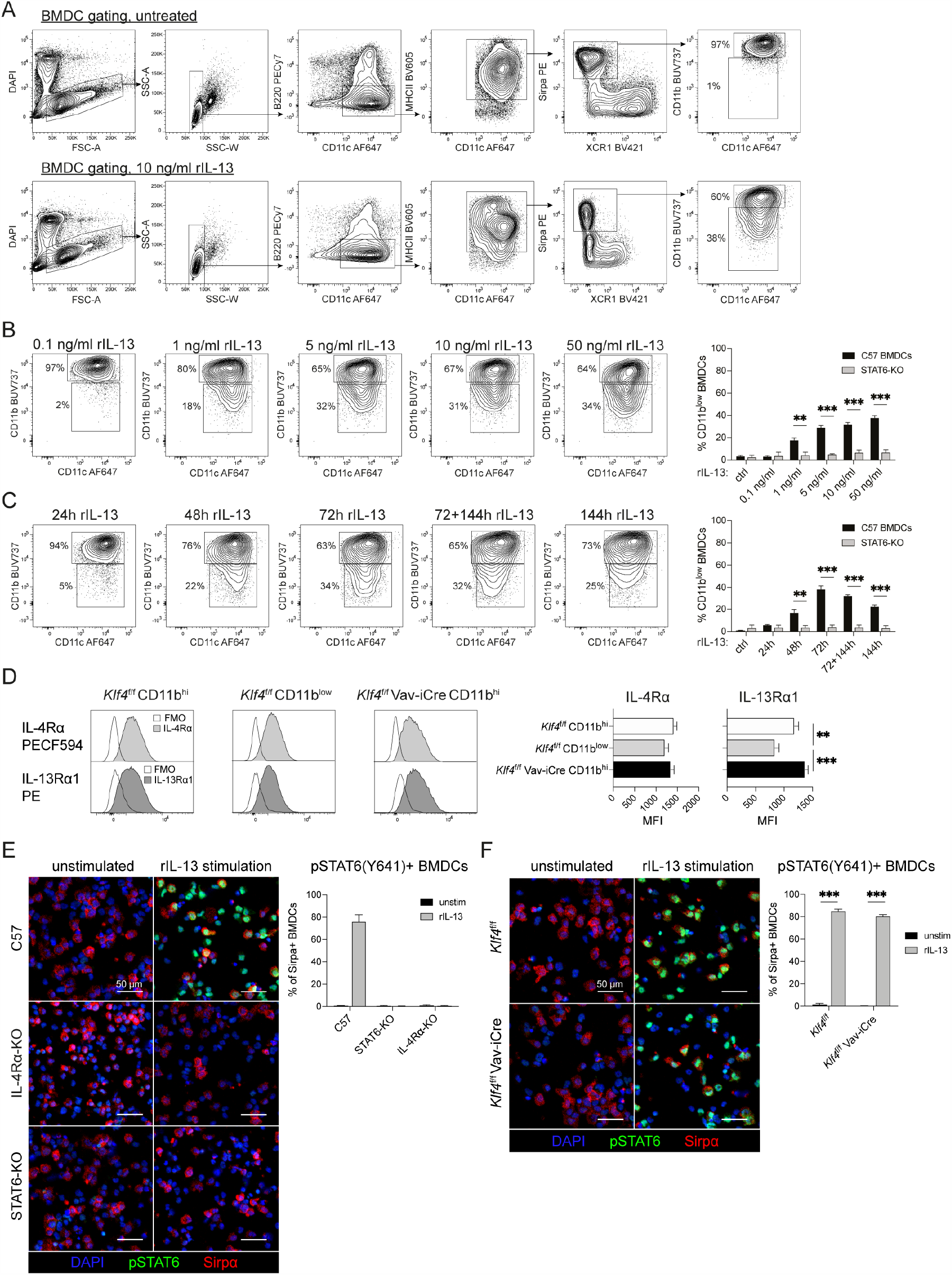
Relating to Figure 4: IL-13 signalling drives the development of CD11b^low^ DC2 *in vitro* in a dose- and time-dependent manner. **(A)** Gating strategy to identify Sirpa+ DC2 in FLT3L BMDC cultures established from C57BL/6 (C57) BM. Cultures were untreated (upper panels) or supplemented with 10 ng/ml rIL-13 for the last 72h of culture (lower panels). **(B)** Representative contour plots showing the phenotype and relative frequencies of Sirpa^+^ DC2 subsets in FLT3L BMDCs cultures from C57 BM. Cells were cultured with the indicated concentrations of rIL-13 for the last 72h of culture. The bar graph shows mean ± SEM, each bar refers to 4 cultures pooled from 2 independent experiments. *P* values were determined using a two-tailed Student’s t-test. **p < 0.01; ***p < 0.001; only significant comparisons are indicated. **(C)** Representative contour plots showing the phenotype and relative frequencies of Sirpa^+^ DC2 subsets in FLT3L BMDC cultures from C57 BM. Cell cultures were supplemented with 10ng/ml rIL-13 at the indicated times before harvest. The bar graph shows mean ± SEM, each bar refers to 4 cultures pooled from 2 independent experiments. *P* values were determined using a two-tailed Student’s t-test. **p < 0.01; ***p < 0.001; only significant comparisons are indicated. **(D)** Representative histograms showing IL-4Rα and IL-13Rα1 expression on DC2 subsets from the skin-draining lymph nodes of *Klf4*^f/f^ → C57 and *Klf4*^f/f^ Vav1-iCre → C57 chimeric mice as determined by fluorescent staining and flow cytometry. Empty histograms refer to FMOs. Histograms show concatenated data from 3 mice. Bar graphs show mean ± SEM for 10-11 mice pooled from 3 independent experiments. *P* values were determined using an ordinary one-way ANOVA with Sidak correction. **p < 0.01; ***p < 0.001; only significant comparisons are indicated. (**E, F**) Representative images of male C57, IL-4Rα-KO, STAT6-KO (E) *Klf4*^f/f^ → C57 and *Klf4*^f/f^ Vav1-iCre → C57 (F) FLT3L BMDCs that were either unstimulated or treated with 100ng/ml rIL-13 for 30 minutes, centrifuged on a glass slide, fixed and stained with anti-Sirpα, anti-phospho-STAT6 (Y641) and DAPI. Scale bars correspond to 50 µm. An average of 2200 Sirpα+ cells/sample were assessed. Bar graphs show the mean frequency of pSTAT6+ cells in the Sirpα+ BMDC population ± range (E) or ± SEM (F), each bar refers to 2 (E) or 5-6 (F) cultures pooled from 2 separate experiments. *P* values were determined using multiple t-tests. ***p < 0.001; only significant comparisons are indicated.

**Figure S5.**
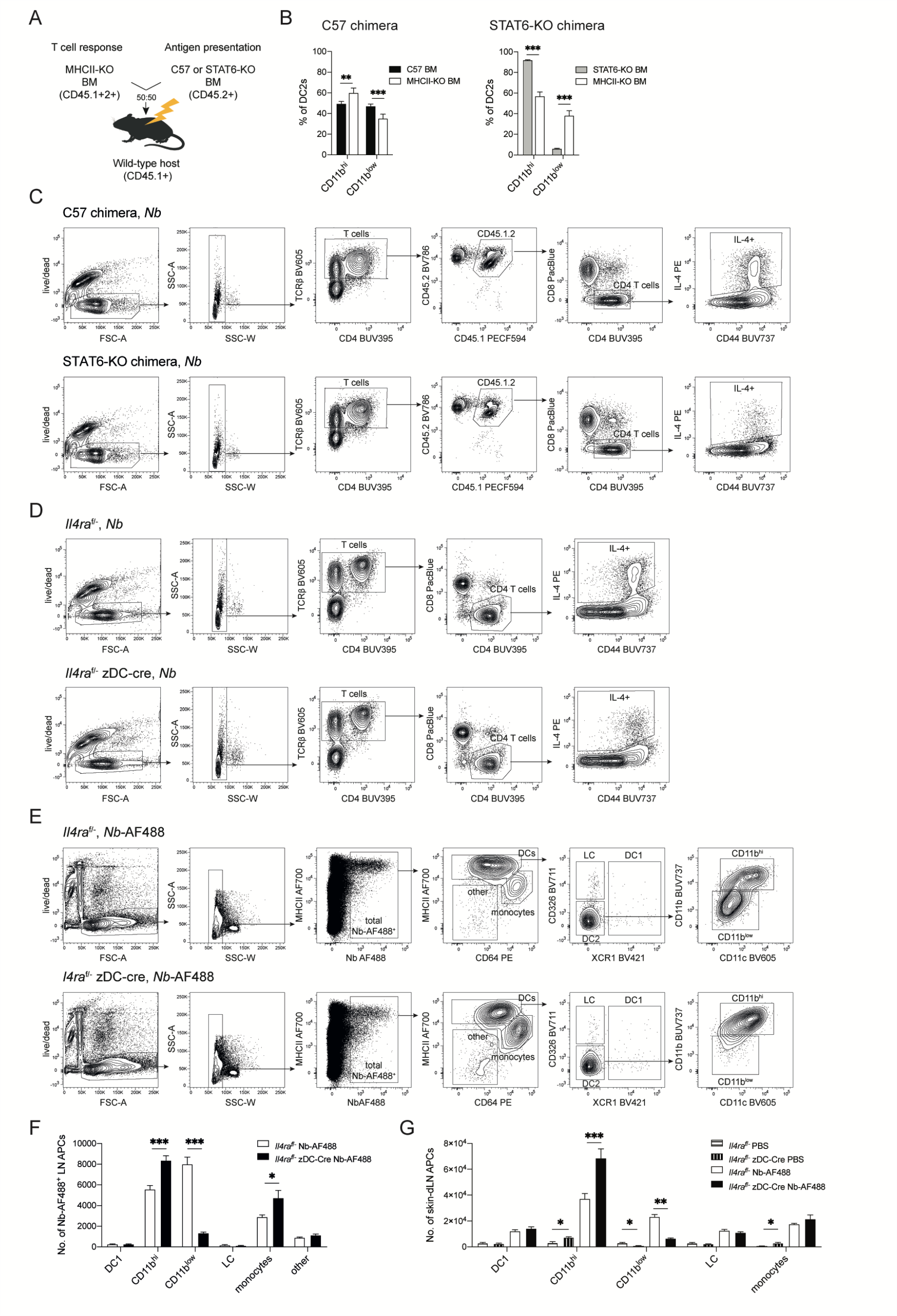
Relating to Figure 5: IL-13 signalling in DC2s is required for optimal IL-4^+^ Th responses in skin-draining lymph node (dLN). (A) Experimental set-up of mixed bone marrow (BM) chimeras in which the antigen-presenting function of MHCII^+^ CD45.2^+^ conventional DCs from either C57 or STAT6-KO BM was assessed by measuring the response of STAT6-sufficient CD45.1^+^2^+^ T cells from MHCII-KO BM. (B) Relative frequencies of MHCII^+^CD45.2^+^ and MHCII-KO CD45.1^+^2^+^ DC2 subsets in the skin-dLN of naïve C57 or STAT6-KO mixed BM chimeras. Bar graphs show mean ± SEM, each bar refers to 8 mice pooled from 2 independent experiments. *P* values were determined using a two-tailed Student’s t-test. **p < 0.01; ***p < 0.001. (C) Gating strategy for cytokine-positive CD45.1^+^2^+^ CD4 T cells from the skin-dLN of C57 or STAT6-KO mixed BM chimeras 5 days after intradermal immunization. Intracelluar cytokine staining was performed after 5h PMA+ionomycin stimulation. Gating for IL-4+ CD4 T cells after *Nb* immunization in C57 (upper panels) or STAT6-KO (lower panels) mixed BM chimeras is shown, other cytokines were gated in a similar manner. (D) Gating strategy for cytokine-positive CD4 T cells from the skin-dLN of *Il4ra*^f/-^ or *Il4ra*^f/-^ zDC-Cre mice 5 days after intradermal immunization. Intracelluar cytokine staining was performed after 5h PMA+ionomycin stimulation. Gating for IL-4+ CD4 T cells after *Nb* immunization in *Il4ra*^f/-^ (upper panels) or *Il4ra*^f/-^ zDC-Cre (lower panels) mice is shown, other cytokines were gated in a similar manner. (E) Gating strategy for AF488^+^ cells in the skin-dLN of *Il4ra*^f/-^ (upper panels) or *Il4ra*^f/-^ zDC-Cre (lower panels) mice 48 hours after intradermal injection of AF488-labelled *Nb* (*Nb*-AF488). (F) Number of AF488^+^ cells in the skin-dLN of *Il4ra*^f/-^ or *Il4ra*^f/-^ zDC-Cre mice 48 hours after intradermal injection of AF488-labelled *Nb*. Bar graphs show mean ± SEM, each bar refers to 7 mice pooled from 2 independent experiments. *P* values were determined using a two-tailed Student’s t-test. *p < 0.05; ***p < 0.001; only significant comparisons are indicated. (G) Number of antigen-presenting cells (APCs) by population in the skin-dLN of *Il4ra*^f/-^ or *Il4ra*^f/-^ zDC-Cre mice 48 hours after intradermal injection of AF488-labelled *Nb* (*Nb*-AF488) or PBS. Bar graphs show mean ± SEM, each bar refers to 3-9 mice from 2 independent experiments. *P* values were determined using a two-tailed Student’s t-test to compare similarly treated groups of different genotype. *p < 0.05; **p < 0.01; ***p < 0.001; only significant comparisons are indicated.

**Figure S6.**
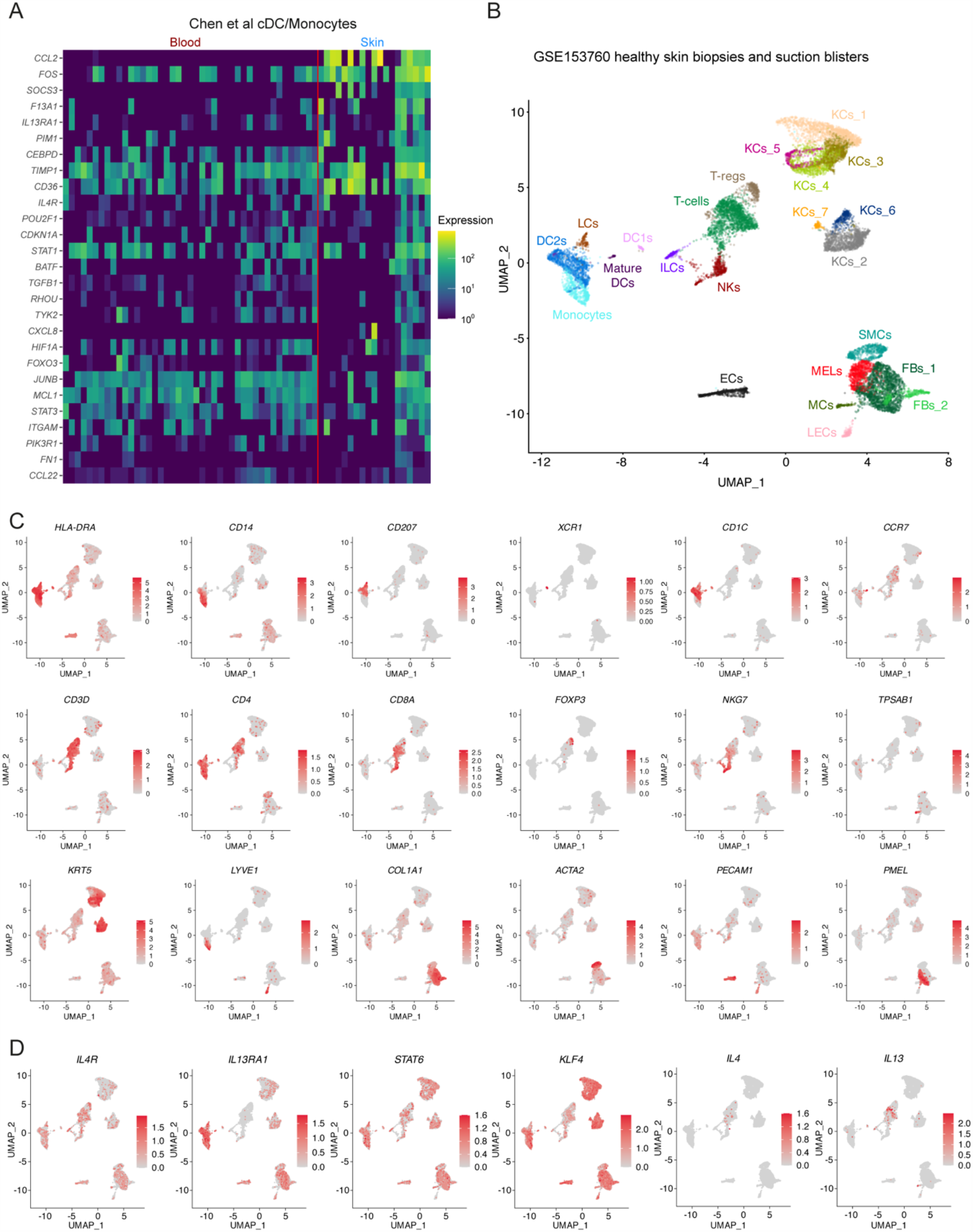
Relating to Figure 6: Enrichment of IL-4/IL-13 signature genes in DC2s from healthy human skin. (A) Heatmap showing the expression of GSEA core-enrichment genes of the IL-4 and IL-13 Reactome pathway in healthy control cDCs and monocytes from blood vs skin as determined by scRNAseq (Chen et al, 2020). (B) UMAP plot showing scRNA-seq subclusters of cells from skin biopsies and suction blisters of healthy controls from the published dataset GSE153760. EC: Endothelial cells, FB_1-2: Fibroblast clusters 1&2, ILC: Innate lymphoid cells, LCs: Langerhans cells, LEC: Lymphendothelial cells, MC: Mast cells, MEL: Melanocytes, NK: Natural killer cells, KC_1-7: Keratinocyte clusters 1-7, SMC: Smooth muscle cells, T-regs: Regulatory T cells. (C) Feature plots of the UMAP clusters in B showing the expression level of cluster-specific transcripts used for cluster identification. Color intensity represents the level of normalized gene expression. (D) Feature plots of the UMAP clusters in B showing the expression levels of *IL4R, IL13RA1, STAT6, KLF4, IL4* and *IL13* transcripts. Color intensity represents the level of normalized gene expression.

